# Extravascular coagulation stabilizes pro-fibrotic stromal states via tumor-intrinsic PAR1 signaling in pancreatic ductal adenocarcinoma

**DOI:** 10.64898/2026.06.25.734662

**Authors:** Sae Rome Choi, Natalia Ospina Munoz, Hye-ran Moon, Sagar M Utturkar, Duy C.K. Do, Yun Chang, Xiaoping Bao, Abigail D. Cox, Timothy L. Ratliff, Claudius Conrad, Melissa L. Fishel, Matthew J. Flick, Nadia A Lanman, Bennett D. Elzey, Bumsoo Han

## Abstract

Pancreatic ductal adenocarcinoma (PDAC) exhibits a desmoplastic stroma with context-dependent tumor-restraining and tumor-promoting functions, highlighting the need to selectively reprogram stromal states. Extravascular coagulation is a prominent feature of the PDAC tumor microenvironment, yet whether it functions as an upstream regulator of fibrotic stromal states, rather than merely a byproduct of tumor-associated vascular dysfunction, has remained unclear. Here, we identify extravascular coagulation as a tumor-amplified regulatory module that stabilizes pro-fibrotic stromal states via tumor-intrinsic protease-activated receptor-1 (PAR1) signaling. To interrogate this axis mechanistically, we integrated human tumor bioinformatics with microphysiological tumor-stroma (MPTS) models that reconstruct tumor-stroma interactions under controlled coagulation exposure, followed by cross-scale validation in vivo. Analysis of The Cancer Genome Atlas (TCGA) revealed heterogeneous F2R (PAR1) expression across tumors, with elevated expression associated with fibrotic transcriptional programs and reduced survival. Consistently, thrombin induced coordinated pro-fibrotic programs in tumor cells and cancer-associated fibroblasts (CAFs), which were recapitulated in MPTS where tumor-intrinsic PAR1 was required for amplification of extracellular matrix deposition and CAF activation. Mechanistically, PAR1 signaling amplified tumor-stroma communication, in part through induction of TGF-β1-dependent pathways, establishing a reinforcing feedback loop that stabilizes fibrotic remodeling. Pharmacologic inhibition of PAR1 selectively suppressed the fibrotic transcriptional program within myofibroblastic CAFs while reducing the abundance of other CAF subtypes, reprogramming stromal states and attenuating tumor progression across MPTS and in vivo models. These findings establish a coagulation-PAR1 axis as an upstream organizer of PDAC stromal architecture and identify pharmacologic PAR1 inhibition as a mechanistically grounded strategy for selectively reprogramming the tumor-promoting stroma.

## INTRODUCTION

Pancreatic ductal adenocarcinoma (PDAC) is one of the most lethal malignancies, with a five-year survival rate below 13% that has improved only marginally over the past two decades [1]. A defining feature of PDAC is its highly desmoplastic tumor microenvironment (TME), in which dense extracellular matrix (ECM) and activated cancer-associated fibroblasts (CAFs) promote therapeutic resistance, immune exclusion, and tumor progression [2]. The recognition that the desmoplastic stroma exerts both tumor-restraining and tumor-promoting functions has complicated efforts to therapeutically target it [3–8]. Strategies aimed at broad stromal depletion, including inhibition of the Sonic Hedgehog pathway (IPI-926/vismodegib) [9, 10], enzymatic degradation of hyaluronan (PEGPH20) [11, 12], and focal adhesion kinase inhibition [13, 14], have each failed to demonstrate clear clinical benefit as monotherapy stromal-targeting strategies, and preclinical studies have demonstrated that indiscriminate CAF depletion can paradoxically accelerate tumor progression and worsen survival [3, 5]. These failures underscore a critical need: rather than eliminating the stroma, effective therapeutic strategies must identify and selectively target the upstream signals that establish and maintain pro-fibrotic, tumor-promoting stromal states while preserving tumor-restraining functions.

Extravascular coagulation is a prominent and clinically well-recognized feature of PDAC. Patients with pancreatic cancer exhibit one of the highest rates of venous thromboembolism among solid tumors, a hypercoagulable phenotype historically described as Trousseau syndrome and now understood to carry independent prognostic significance [15, 16]. This systemic phenotype has a local counterpart within the TME: leaky tumor vasculature [17–19] permits circulating coagulation factors to enter the interstitial space, where tissue factor (TF) expressed by pancreatic cancer cells (PCCs) initiates local coagulation cascade activation and generates thrombin [20]. Beyond its canonical role in fibrin deposition, thrombin engages cellular signaling through protease-activated receptors (PARs), particularly PAR1, which is expressed across tumor cells, CAFs, endothelial cells, and immune cells and has established roles in tumor cell proliferation, invasion, and immune evasion [21–25]. These observations position extravascular coagulation not merely as a systemic complication of tumor progression, but as an active signaling input into the TME.

Despite this body of work, whether extravascular coagulation functions as an upstream regulator of fibrosis-associated stromal states in PDAC remains unknown. Prior studies of thrombin-PAR1 signaling in PDAC have established its roles in PCC proliferation, immune evasion, and chemoresistance [22–25]. These effects are attributable predominantly to PAR1 activity within the tumor cell compartment. Whether tumor-intrinsic PAR1 signaling extends its influence to the stromal compartment, driving CAF activation and the establishment of pro-fibrotic ECM programs, has not been examined. This distinction is mechanistically important: if coagulation acts as an upstream organizer of stromal state architecture through a tumor-CAF paracrine circuit, it would define a regulatory axis that is both upstream of and distinct from the individual stromal components (TGF-β, ECM, CAF subpopulations) that have been targeted downstream. Given that indiscriminate stromal depletion can render PDAC more aggressive [3, 5], a strategy that reprograms fibrosis-associated stromal states at their upstream regulatory source, rather than eliminating stromal components outright, may avoid this risk while still disrupting tumor-promoting stromal function. Resolving whether coagulation-PAR1 signaling serves as such a control node would provide a mechanistic basis for selectively reprogramming the PDAC stroma, rather than disrupting its individual downstream outputs.

Here, we address this gap using an integrated mechanistic and therapeutic analysis spanning human tumor bioinformatics, microphysiological tumor-stroma models, and in vivo validation. We developed a microphysiological tumor-stroma (MPTS) model that reconstitutes extravascular coagulation within a controlled PDAC microenvironment, enabling cell type-specific perturbation of PAR1 signaling. We hypothesized that extravascular coagulation, acting through tumor-intrinsic thrombin-PAR1 signaling, functions as an upstream driver of fibrosis-associated stromal states in PDAC. To test this, we first examined whether coagulation and PAR1 pathway activity associate with fibrotic transcriptional programs and clinical outcome in human PDAC tumors. We then used the MPTS platform to determine whether thrombin-PAR1 signaling is sufficient to drive tumor-derived paracrine signaling, CAF activation, and ECM remodeling, and to identify the downstream mediator linking tumor-intrinsic PAR1 activation to stromal responses. Finally, we tested whether pharmacologic PAR1 inhibition could reverse this axis, evaluating its effects on stromal reprogramming and tumor progression in both microphysiological and in vivo PDAC models.

## MATERIALS AND METHODS

### Cell Culture and Reagents

Murine pancreatic cancer (KPC2) and cancer-associated fibroblast (mCAF) cell lines were isolated from genetically engineered KPC mice (Kras^G12D/+^; p53^R172H/+^; Elas^CreER/+^). Human PDAC cell lines (Panc10.05, Panc1, MIA PaCa-2) and CAF lines (CAF19, 02-hT) were cultured under standard conditions in RPMI or DMEM/F12 supplemented with 2.05 mM L-glutamine, 100 μg/mL penicillin/streptomycin, and 5% or 10% FBS. Cells were maintained below 80% confluence and used within defined passage limits. Cryopreservation was performed using 90% FBS and 10% DMSO. Tumor and CAF cells were fluorescently labeled with TdTomato and GFP, respectively.

### RT-qPCR

Gene expression was quantified by RT-qPCR following RNA isolation and cDNA synthesis. Expression levels of genes of interest were normalized to 18S RNA using the ΔΔCt method. Primer and probe sequences are provided in Supplementary Methods.

### RNA-seq and transcriptomic analysis

Total RNA was isolated from KPC tumor cells and mCAFs following thrombin stimulation under defined experimental conditions. Cells were serum-starved (1% FBS, 24 h) prior to stimulation with thrombin (1 U/mL, 24 h; n = 3 per group). RNA samples were quality-checked and sequenced using paired-end Illumina platforms with a minimum depth of 30 million reads per sample. Raw sequencing reads were subjected to quality control, including adapter trimming and filtering based on Phred score and read length. High-quality reads were aligned to the mouse reference genome (GRCm38) using splice-aware aligners. Gene-level counts were generated using feature-based quantification tools. Differential expression analysis was performed using the edgeR framework with multiple testing correction. Genes meeting false discovery rate (FDR) thresholds were considered differentially expressed. Pathway enrichment analysis was conducted using Ingenuity Pathway Analysis (IPA) and Gene Set Enrichment Analysis (GSEA) with MSigDB gene sets to identify thrombin-PAR1-associated transcriptional programs. Co-expressed gene clusters were identified, and heatmaps were generated to visualize expression patterns across conditions. Detailed procedures are described in Supplementary Methods.

### Human dataset analysis

The Cancer Genome Atlas (TCGA) Pan-Cancer Atlas dataset (cBioPortal study ID: paad_tcga_pan_can_atlas_2018) was analyzed, which primarily consists of resected primary tumor samples. The dataset initially encompassed 184 patients (101 male, 83 female). For each patient, raw RNA-Seq by Expectation-Maximization (RSEM) expression values were extracted. Any patients with missing expression data, types of normal or metastasized tissues were excluded from the analysis. To normalize the data and minimize outlier skewness, raw values were log-transformed and converted to z-scores, resulting in a distribution with a mean of approximately zero. Samples were then stratified into three expression groups based on quartiles: the Low group (bottom 25%), the High group (top 25%), and the Medium group (interquartile range) for F2R and TGFB1 expression. All statistical evaluations were performed using OriginPro 2026 (OriginLab Corporation, Northampton, MA, USA). Qualitative variables were compared across groups using chi-square tests. Kaplan-Meier (K-M) methods were utilized to estimate the probability of Disease-Specific Survival (DSS) over a 60-month period. Differences in survival trajectories among the three expression groups were evaluated, and a log-rank p-value was calculated to determine statistical significance. Hazard ratios (HR) were estimated utilizing a Cox proportional-hazards model. To identify the biological pathways and functional signatures associated with F2R expression, GSEA was conducted.

### Microphysiological tumor-stroma (MPTS) model

Microfluidic MPTS devices were fabricated using soft lithography, as described previously [26–28], to reconstruct fibrosis-associated tumor-stroma signaling under controlled coagulation conditions in a 3D tumor microenvironment. Tumor cells and CAFs were embedded in type I collagen and exposed to thrombin and/or PAR1 inhibitor (vorapaxar) under controlled flow conditions. Detailed fabrication and culture procedures are described in Supplementary Methods.

### Quantification of phenotypes

Tumor-stroma phenotypes were assessed using fluorescence imaging, ELISA, collagen secretion assays, and immunostaining. Quantitative image analysis was performed using ImageJ-based segmentation methods. Detailed protocols are provided in Supplementary Methods.

### Spatial transcriptomics and treatment

Orthotopic KPC tumor-bearing mice were treated with vorapaxar (VPX) or vehicle control under approved institutional protocols. Tumors were harvested at endpoint, flash-frozen, and processed for spatial transcriptomic analysis. Spatial gene expression profiling was performed using the 10x Genomics Visium platform on fresh-frozen tumor sections. Tissue sections were processed according to the manufacturer’s protocol, including permeabilization, cDNA synthesis, library preparation, and sequencing. Sequencing data were processed using the Space Ranger pipeline to generate spatially resolved gene expression matrices. Cell-type annotation and spatial clustering were performed using reference transcriptomic datasets and marker gene-based curation. Differential expression analysis was conducted to identify transcriptional changes across spatial regions and treatment conditions. Detailed protocols are provided in Supplementary Methods.

### Xenograft model and treatment

Human PCC-CAF xenografts were established in NRG mice using co-injection of tumor and CAF cells. MIA PaCa-2 and CAF19 cells were injected subcutaneously (1:5 ratio, 0.7×10□ : 3.5×10□ cells) into NRG mice in PBS:Matrigel (1:1). After 14 days, VPX (0–20 mg/kg) was administered via oral gavage using a 5-day-on/2-day-off schedule for 2 weeks. Tumor growth was monitored and tissues collected at Day 14 for analysis.

### Histology and immunofluorescence

For in vitro samples, cells were fixed with 4% formaldehyde, permeabilized with Triton-X, and stained for αSMA using primary and biotinylated secondary antibodies. Detection was performed using streptavidin-Alexa Fluor conjugates. Imaging was performed using confocal microscopy.

For tissue samples, tumors were fixed in formalin, embedded in paraffin, and sectioned. H&E staining was performed using automated systems. For immunofluorescence, sections underwent antigen retrieval, blocking, and staining with antibodies against collagen, RFP, GFP, and αSMA. Quantification was performed using ImageJ across multiple regions of interest.

### Statistical analysis

Data are presented as mean ± standard error (N ≥ 3). Statistical significance was determined using Student’s t-test or one-way ANOVA with post hoc Tukey test, with p ≤ 0.05 considered significant. Error bars represent standard error.

## RESULTS

### Coagulation-fibrosis coupling defines a poor-prognosis stromal state in PDAC

To assess whether coagulation signaling is active in human PDAC, we examined PAR1 and TF expression in pancreatic tumors. Immunofluorescence analysis revealed that both proteins were strongly upregulated in tumor tissues (**Fig. 1A**), accompanied by elevated expression of the fibrosis marker α-SMA. In contrast, normal adjacent pancreas exhibited minimal expression of TF, PAR1, and α-SMA, consistent with activation of extravascular coagulation and desmoplastic signaling in PDAC. Additional tissue samples are presented in the supporting information (**Fig. S1**).

**Figure 1.**
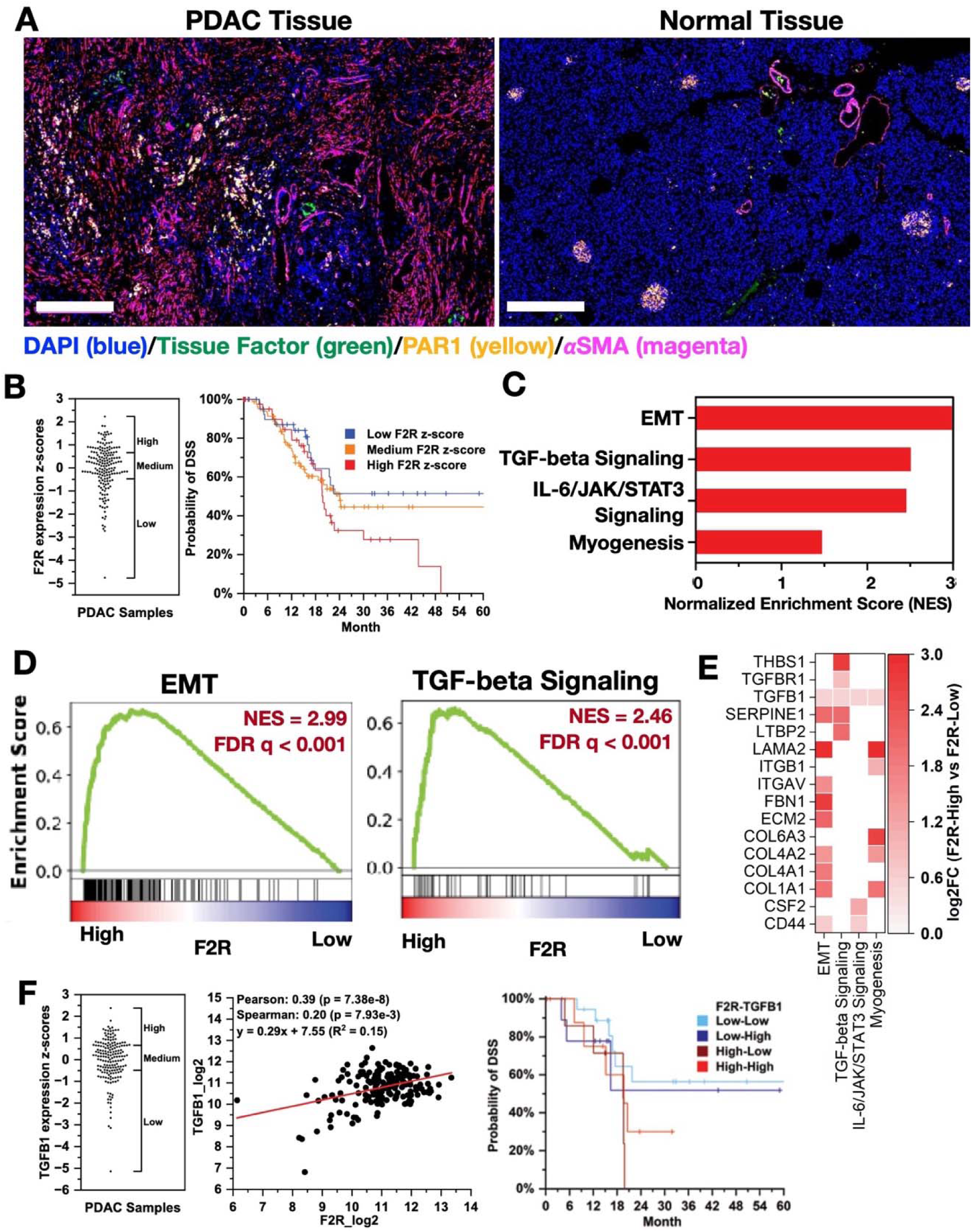
Coagulation signaling and fibrotic transcriptional programs in human PDAC. (A) Immunofluorescence staining of tissue factor (green), PAR1 (yellow), and α-SMA (magenta) in PDAC tumor and normal pancreatic tissue. Nuclei are stained with DAPI (blue). Scale bar = 300 μm. (B) Distribution of F2R expression z-scores across The Cancer Genome Atlas (TCGA) PDAC cohort (left) and Kaplan-Meier analysis of disease-specific survival (DSS) stratified by F2R expression (right). (C) Normalized enrichment scores for curated fibrosis-associated Hallmark gene sets in F2R-high versus F2R-low tumors. (D) Running enrichment plots for the epithelial-mesenchymal transition and TGF-β signaling gene sets. (E) Heat map of leading-edge genes across the enriched gene sets, shown as log2 fold-change (F2R-high versus F2R-low). TGFB1 was included based on its role in the study hypothesis and did not rank among the most highly differentially expressed leading-edge genes. (F) Distribution of TGFB1 expression z-scores (left), correlation between F2R and TGFB1 expression (middle), and Kaplan-Meier analysis of DSS stratified by combined F2R and TGFB1 expression (right).

Analysis of The Cancer Genome Atlas (TCGA) PDAC cohort revealed heterogeneous expression of F2R, encoding PAR1, across tumors (**Fig. 1B**). Patients were stratified by F2R expression z-score into high, medium, and low groups. Kaplan-Meier analysis demonstrated that patients with F2R-high tumors exhibited reduced disease-specific survival (DSS) compared with the F2R-low group, which maintained approximately 50% survival up to 60 months. Patients with intermediate F2R expression displayed survival between these two groups, establishing a graded relationship between F2R expression and disease progression. The association between elevated F2R expression and poor survival suggests that coagulation-dependent stromal states may contribute to clinically aggressive disease phenotypes.

To explore the transcriptional programs associated with elevated PAR1 signaling, gene set enrichment analysis (GSEA) was performed comparing F2R-high and F2R-low tumors against the Hallmark collection, and pathways relevant to fibrotic stromal remodeling were examined (**Fig. 1C**). Epithelial-mesenchymal transition (NES = 2.99), TGF-β signaling (NES = 2.46), IL-6/JAK/STAT3 signaling (NES = 2.44), and myogenesis (NES = 1.47) were enriched in F2R-high tumors. Running enrichment plots for the epithelial-mesenchymal transition and TGF-β signaling gene sets are shown in **Fig. 1D** (FDR q < 0.001 for both). The complete Hallmark analysis is presented in the supporting information (**Fig. S2**).

Leading-edge analysis across these enriched gene sets identified a shared set of extracellular matrix and fibrosis-associated genes elevated in F2R-high tumors, including THBS1, SERPINE1, LTBP2, FBN1, LAMA2, ECM2, and multiple collagen genes (COL1A1, COL4A1, COL4A2, COL6A3), together with integrins (ITGAV, ITGB1) and TGF-β pathway components (TGFB1, TGFBR1) (**Fig. 1E**). Several of these genes, including THBS1, SERPINE1, and LTBP2, are associated with the storage and activation of latent TGF-β, suggesting that TGF-β pathway activity in F2R-high tumors may be regulated at multiple levels. TGFB1 itself showed a comparatively modest fold-change but was represented across the enriched gene sets, consistent with a broadly distributed rather than a strongly amplitude-driven contribution.

Given the recurrence of TGF-β pathway components across these programs, we next examined TGFB1 directly, a key mediator of fibrotic signaling [29–31]. TGFB1 expression was similarly heterogeneous across the cohort and showed a modest but significant positive correlation with F2R expression (Pearson r = 0.39; Spearman ρ = 0.20), indicating transcriptional coupling between coagulation signaling and fibrotic programs. Patients were then co-stratified by F2R and TGFB1 expression (**Fig. 1F**). Survival patterns were largely dictated by F2R status, with the F2R-high/TGFB1-high group showing the poorest outcome, while TGFB1 status did not clearly separate survival within either F2R stratum. These correlative findings raise the possibility that PAR1 signaling acts as an upstream determinant of fibrotic stromal state and clinical outcome, with TGF-β1 serving as a downstream amplifier of this program, a hypothesis we address directly in the murine and human experimental systems described below.

### An engineered microphysiological tumor-stroma platform enables compartment-resolved interrogation of the tumor-stroma-coagulation axis

Based on these observations, we hypothesized that thrombin-PAR1 signaling amplifies tumor-stroma communication within a tumor-stroma-coagulation axis and contributes to desmoplastic remodeling of the TME. Specifically, thrombin is proposed to activate PAR1 in both PCCs and CAFs, promoting CAF proliferation, extracellular matrix deposition, and CAF-mediated contractility. We further posited that PAR1 suppression may reprogram the PDAC stroma without compromising tumor-restraining functions, motivating development of a system enabling compartment-resolved perturbation of PAR1 signaling.

To enable mechanistic interrogation of this axis, we developed an MPTS model that recapitulates extravascular coagulation within a controlled PDAC microenvironment (**Fig. 2A**). In this system, thrombin introduced through the perfusion channel mimics its accumulation in the tumor interstitium and establishes controlled exposure across the collagen type I matrix, enabling tumor and stromal compartments to experience a defined coagulation-associated microenvironment. Confocal imaging and three-dimensional reconstruction confirmed the formation of integrated PCC-CAF networks within the matrix, with extensive cell-cell contact and alignment (**Fig. 2A**; Supplementary Video 1), consistent with physiologic tumor-stroma architecture.

**Figure 2.**
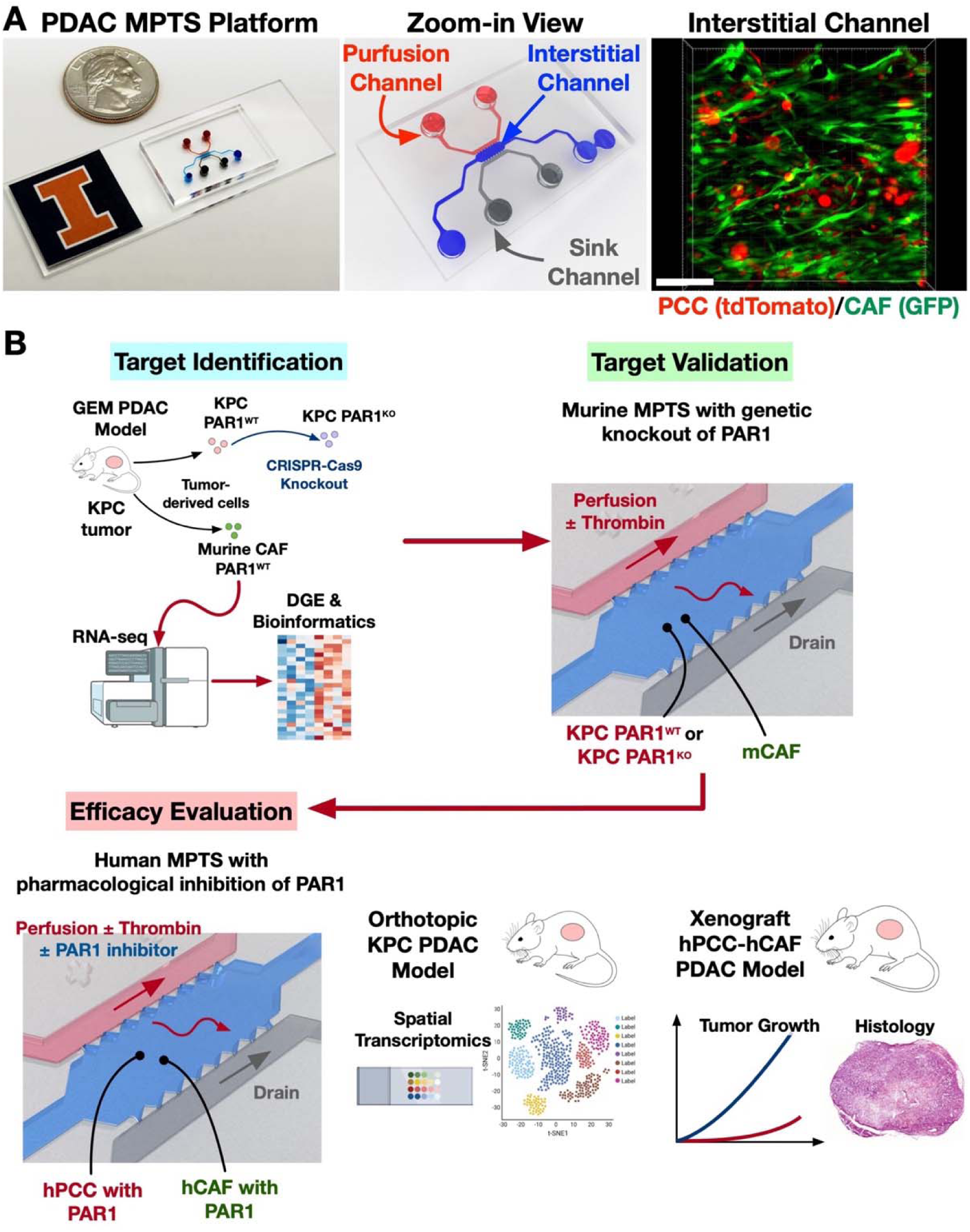
Microphysiological tumor-stroma (MPTS) platform and cross-scale experimental workflow. **(A)** The MPTS platform, shown as a device photograph (left), a schematic of the perfusion, interstitial, and sink channels (middle), and a confocal micrograph with three-dimensional reconstruction of integrated PCC-CAF networks within the interstitial channel (right). Thrombin introduced via the perfusion channel establishes controlled, physiologically relevant exposure to coagulation-dependent stimuli across a collagen type I matrix, reconstituting extravascular coagulation within a PDAC tumor-stroma microenvironment. Scale bar = 100 μm. (B) Cross-scale experimental workflow integrating MPTS models and in vivo systems for systematic target identification, validation, and efficacy evaluation, enabling controlled genetic and pharmacologic perturbation of PAR1 across murine and human contexts.

To investigate whether coagulation signaling contributes to the fibrotic PDAC TME, we implemented a cross-scale translational workflow centered on the tumor-stroma-coagulation axis, integrating MPTS models and in vivo systems for systematic target identification, validation, and efficacy evaluation (**Fig. 2B**). This framework enabled controlled genetic and pharmacologic perturbation of PAR1 to interrogate coagulation-dependent tumor-stroma communication. Tumor-derived KPC cells from Kras^G12D/+^; p53^R172H/+^; Elas^CreER/+^ mice were used to generate PAR1 knockout (KPC PAR1^KO^) lines alongside wild-type controls (KPC PAR1^WT^) as previously reported [22], while murine cancer-associated fibroblasts (mCAFs) enabled modeling of reciprocal tumor-stroma interactions. Human PDAC cell lines were subsequently used to evaluate pharmacologic PAR1 inhibition in MPTS models and to establish xenograft systems for in vivo validation.

### Thrombin induces coordinated pro-fibrotic signaling and tumor-stroma-coagulation coupling in murine systems

To determine whether coagulation signaling modulates tumor and stromal responses, we examined the effect of thrombin on genetically engineered murine PCC and CAF populations - KPC Par1^KO^, KPC Par1^WT^ and mCAFs. Quantitative PCR analysis confirmed that PAR1 expression increased in KPC WT cells following thrombin stimulation, whereas expression was minimal as expected in KPC Par1 KO cells (**Fig. 3A**). Thrombin treatment also induced PAR1 expression in mCAFs, extending this response across both tumor and stromal compartments.

**Figure 3.**
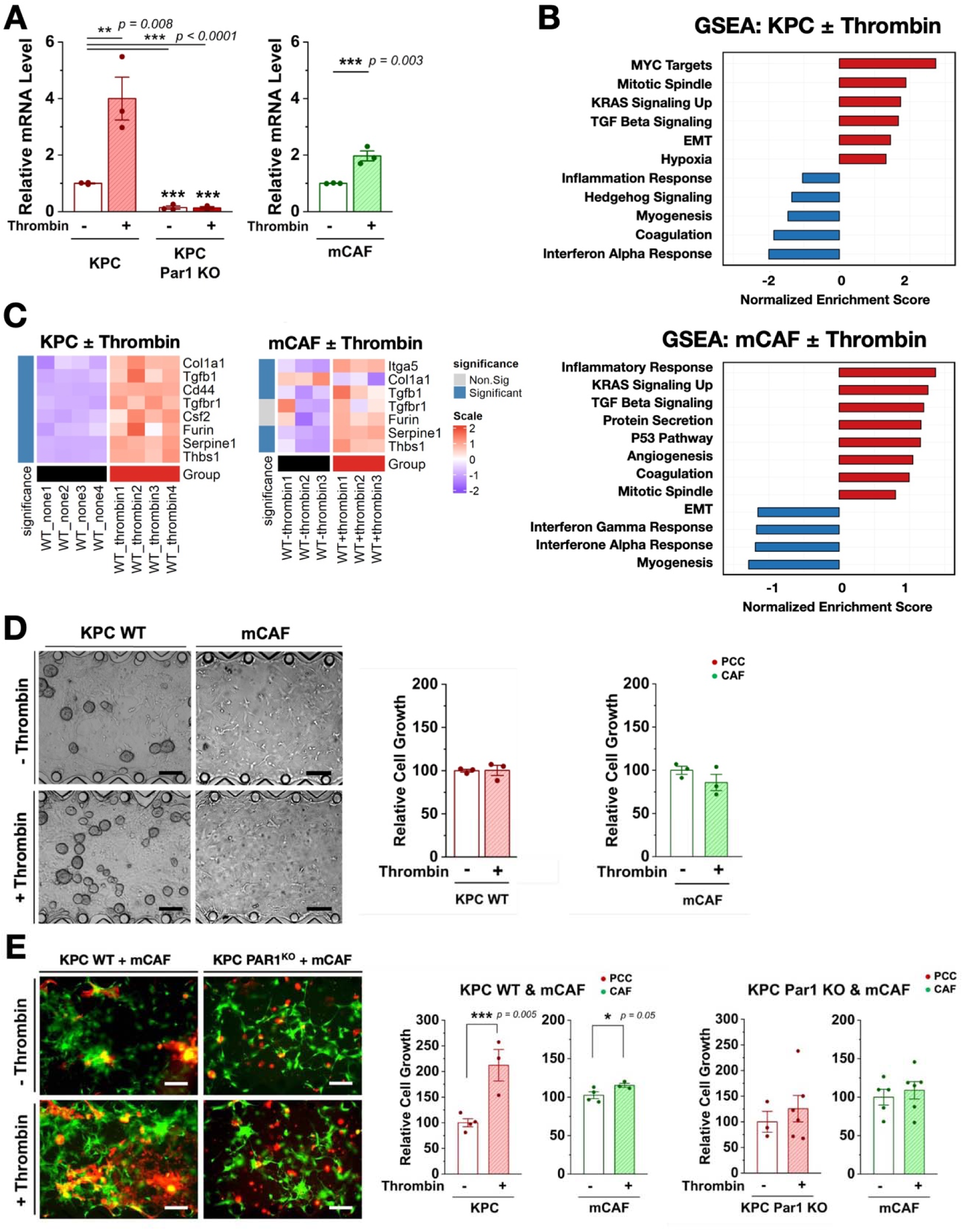
Thrombin-induced transcriptional programs and tumor-stroma interactions in murine systems. (A) Quantitative PCR analysis of PAR1 expression in KPC Par1 WT, KPC Par1 KO, and mCAFs with and without thrombin treatment after 8 days of culture. Statistical analysis with one-way ANOVA with post hoc Tukey test. (B) Gene set enrichment analysis (GSEA) of RNA-seq data from KPC cells and mCAFs following thrombin stimulation. (C) Heatmap of representative differentially expressed genes (DEGs) in KPC cells and mCAFs following thrombin treatment. (D) Representative micrographs and quantification of cell growth in monoculture conditions with and without thrombin treatment. Scale bar = 200 μm. (E) Representative micrographs and quantification of cell growth in co-culture conditions with and without thrombin treatment after 8 days of culture. Scale bar = 200 μm. Statistical analysis with Student’s t-test (N ≥ 3).

We next performed transcriptomic profiling to define thrombin-induced signaling programs. RNA-seq analysis was conducted on KPC Par1^WT^ and mCAF cell lines. GSEA revealed that thrombin stimulation in KPC Par1^WT^ cells enriched KRAS signaling, TGF-β signaling, and epithelial-to-mesenchymal transition (EMT) pathways, consistent with PDAC progression and stromal remodeling (**Fig. 3B**). In parallel, mCAFs exhibited enrichment of inflammatory response, KRAS signaling, TGF-β signaling, angiogenesis, and coagulation pathways, indicating coordinated but compartment-specific transcriptional activation within a fibrosis-associated tumor-stroma context.

To further delineate shared transcriptional responses, we analyzed genes within commonly upregulated pathways, focusing on TGF-β and KRAS signaling. Hallmark genes associated with these pathways are shown in the supplement information (**Fig. S3**). Heatmap analysis of differential gene expression, in conjunction with pathway enrichments, revealed induction of pro-fibrotic and signaling-associated genes across both compartments (**Fig. 3C**). In KPC cells, some of the upregulated genes included *Tgfb1*, *Tgfbr1*, *Thbs1*, *Furin*, *Col1a1*, *Cd44*, *Serpine1*, and *Csf2*, while mCAFs showed increased expression of *Tgfb1*, *Thbs1*, *Itga5*, and *Serpine1*. These transcriptional changes support establishment of a pro-fibrotic stromal signaling state.

The induction of *Tgfb1* across tumor and stromal compartments identified TGF-β1 as a candidate mediator of intercompartmental signaling. Other enriched genes map onto functional modules of fibrotic remodeling and fibrinolysis: *Thbs1* promotes activation of latent TGF-β [32, 33], Furin mediates proteolytic activation of TGF-β and related factors [34–36], and *Serpine1* (expressed as Plasminogen Activator Inhibitor-1, PAI-1) inhibits fibrinolysis and stabilizes fibrin-rich matrices [37–40]. Notably, the lack of significant induction of *Tgfbr1* in CAFs and the reduction of *Col1a1* expression in mCAFs indicate that thrombin-PAR1 signaling alone is insufficient to directly drive canonical fibroblast activation, supporting a model in which tumor-derived signals dominate stromal activation within the tumor-stroma-coagulation axis.

To determine whether these transcriptional programs translate into functional coupling, we next examined thrombin-driven responses using the murine MPTS model where cells were exposed to 0.2 U/mL thrombin for eight days. In monoculture conditions, thrombin stimulation did not significantly alter the growth of KPC Par1^WT^ or mCAFs (**Fig. 3D**). In addition, no difference in growth was observed by Par1^KO^ compared with Par1^WT^ (**Fig. S4**). In co-culture, however, thrombin stimulation enhanced tumor cell growth in Par1^WT^-mCAF conditions, whereas this effect was drastically diminished in KPC Par1^KO^-mCAF co-cultures (**Fig. 3E**), isolating PAR1-dependent tumor-stroma coupling as the dominant driver of the growth response within the tumor-stroma-coagulation axis.

### Tumor-intrinsic PAR1 signaling and heterogeneous stromal responses in human PDAC models

To determine how the tumor-stroma-coagulation axis manifests across heterogeneous human PDAC contexts, we evaluated human PDAC cell lines Panc10.05, Panc1, and MIA PaCa-2 alongside the cancer-associated fibroblast cell line CAF19. The cell lines were selected based on varying levels of TF and PAR1 expression, as determined by qPCR, ranging from low in Panc10.05 to high in MIA PaCa-2 as compared to human pancreatic ductal epithelial cells (HPDE) (**Fig. S5** for TF and **Fig. 4A** for PAR1). This variation enables systematic interrogation of PAR1-dependent tumor-stroma coupling under thrombin stimulation while maintaining CAF19 as a constant stromal component, providing a controlled framework to resolve tumor-intrinsic contributions to stromal activation.

**Figure 4.**
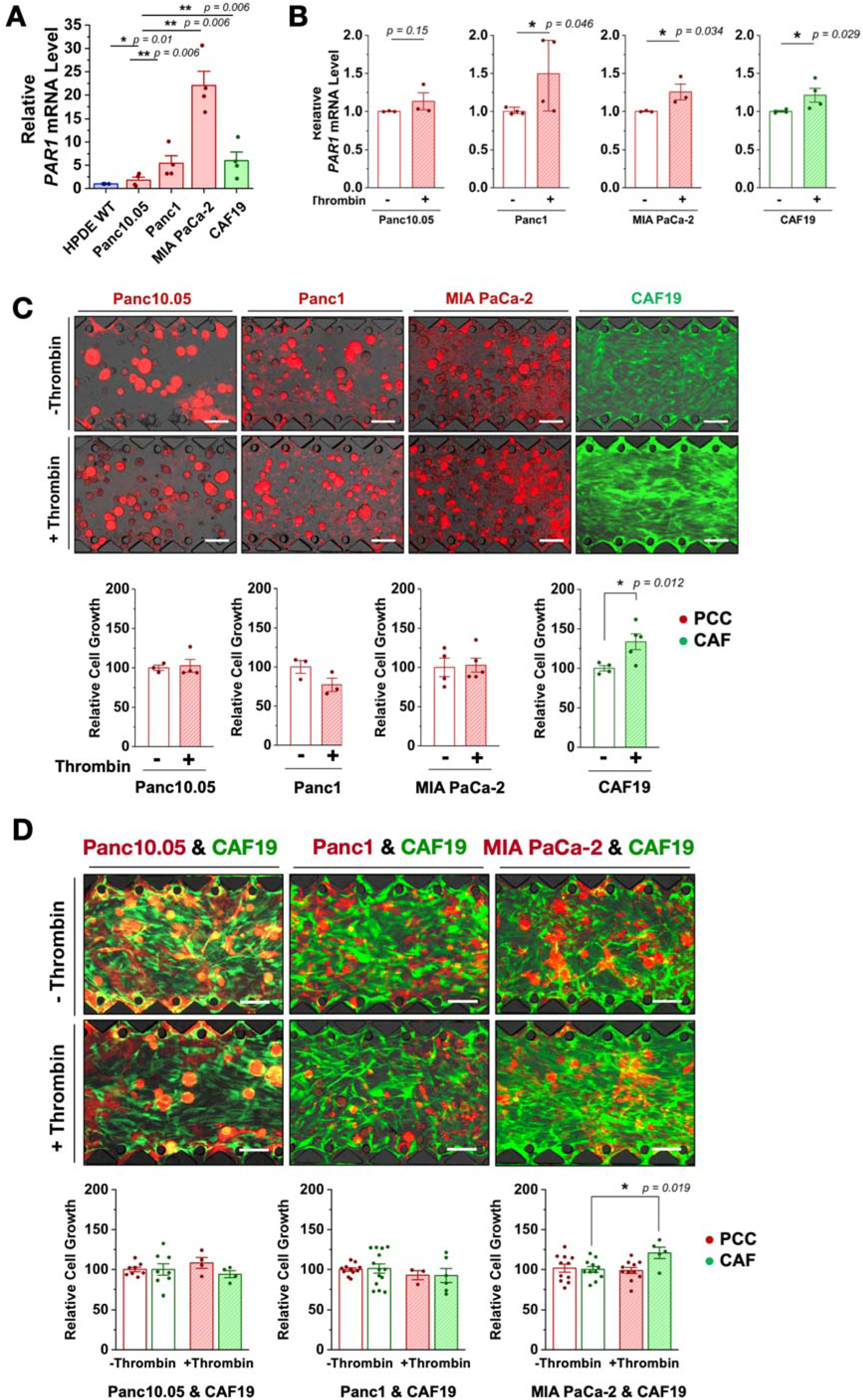
Context-dependent tumor–stroma responses to thrombin in human PDAC models. (A) Quantitative qPCR analysis of PAR1 expression across human PDAC cell lines (Panc10.05, Panc1, MIA PaCa-2) and CAF19. Statistical analysis with one-way ANOVA with post hoc Tukey test. (B) Quantitative qPCR analysis of PAR1 expression in PCCs and CAF19 with and without thrombin treatment. (C) Representative micrographs and quantification of cell growth in monoculture conditions with and without thrombin treatment. (D) Representative micrographs and quantification of cell growth in co-culture conditions with and without thrombin treatment. Scale bar = 200 μm. Statistical analysis with Student’s t test (N ≥ 3).

PAR1 mRNA expression in PCCs generally demonstrated increase in response to thrombin treatment (**Fig. 4B**). These cell lines were introduced into the MPTS platform and continuously exposed to 0.2 U/mL thrombin for eight days. In monoculture, thrombin treatment did not significantly affect the growth or morphology of PDAC cells. In contrast, CAF19 exhibited a notable increase in proliferation under thrombin stimulation, while PDAC cell lines remained largely unchanged (**Fig. 4C**), shifting the dominant response toward the stromal compartment under isolated conditions.

We next evaluated PAR1-dependent tumor-stroma coupling using co-culture models of human PCC-CAF pairs. Thrombin treatment did not significantly increase PCC growth across all tested combinations (**Fig. 4D**). CAF19 proliferation was not uniformly enhanced under co-culture conditions, despite the increase observed in monoculture. A significant increase in CAF growth (∼1.2-fold) was observed only in the MIA PaCa-2-CAF19 co-culture following thrombin treatment, accompanied by a more elongated CAF morphology consistent with an activated state. Given that MIA PaCa-2 cells express the highest levels of PAR1, this response localizes stromal activation to tumor-intrinsic PAR1-high contexts and again suggests that heterogeneity in the tumor leads to differential activation of the stromal compartment.

In contrast to murine systems (**Fig. 3**), where thrombin stimulation promoted robust tumor growth through PAR1-dependent tumor-stroma coupling, human models exhibited selective stromal activation, linked to tumor-intrinsic PAR1 expression. Across these systems, thrombin-PAR1 signaling engaged a conserved tumor-stroma-coagulation axis, while downstream phenotypic outputs varied according to baseline signaling states and tumor context. These findings suggest that tumor proliferation represents only one functional output of the thrombin-PAR1 axis and may not be its predominant manifestation across human PDAC contexts; other fibrosis-associated microenvironmental programs, including ECM remodeling and stromal activation, may instead dominate

### Tumor-derived TGF-β1 mediates PAR1-dependent stromal activation and ECM remodeling

In addition to effects on proliferation through PAR1 signaling, the RNA-seq analysis (**Fig. 3**) identified a suite of genes important in tumor-stroma crosstalk that were upregulated in response to thrombin. To validate these findings in human PCCs and CAFs, we performed RT-qPCR to measure mRNA levels of key genes identified through differential expression analyses, focusing on the MIA PaCa-2–CAF19 pair, which exhibited the most robust response to thrombin (**Fig. 4**). In MIA PaCa-2 cells, thrombin treatment significantly increased expression of fibrosis-associated genes, most prominently TGFB1 (**Fig. 5A**). CSF2 (colony stimulating factor 2) also showed a trend toward increased expression, suggesting a potential role in modulating tumor-stroma and immune interactions. These findings identify TGF-β1 as a major downstream mediator of tumor-intrinsic PAR1 signaling within the tumor-stroma-coagulation axis, with PAR1 functioning as a control node governing tumor-derived TGF-β signaling.

**Figure 5.**
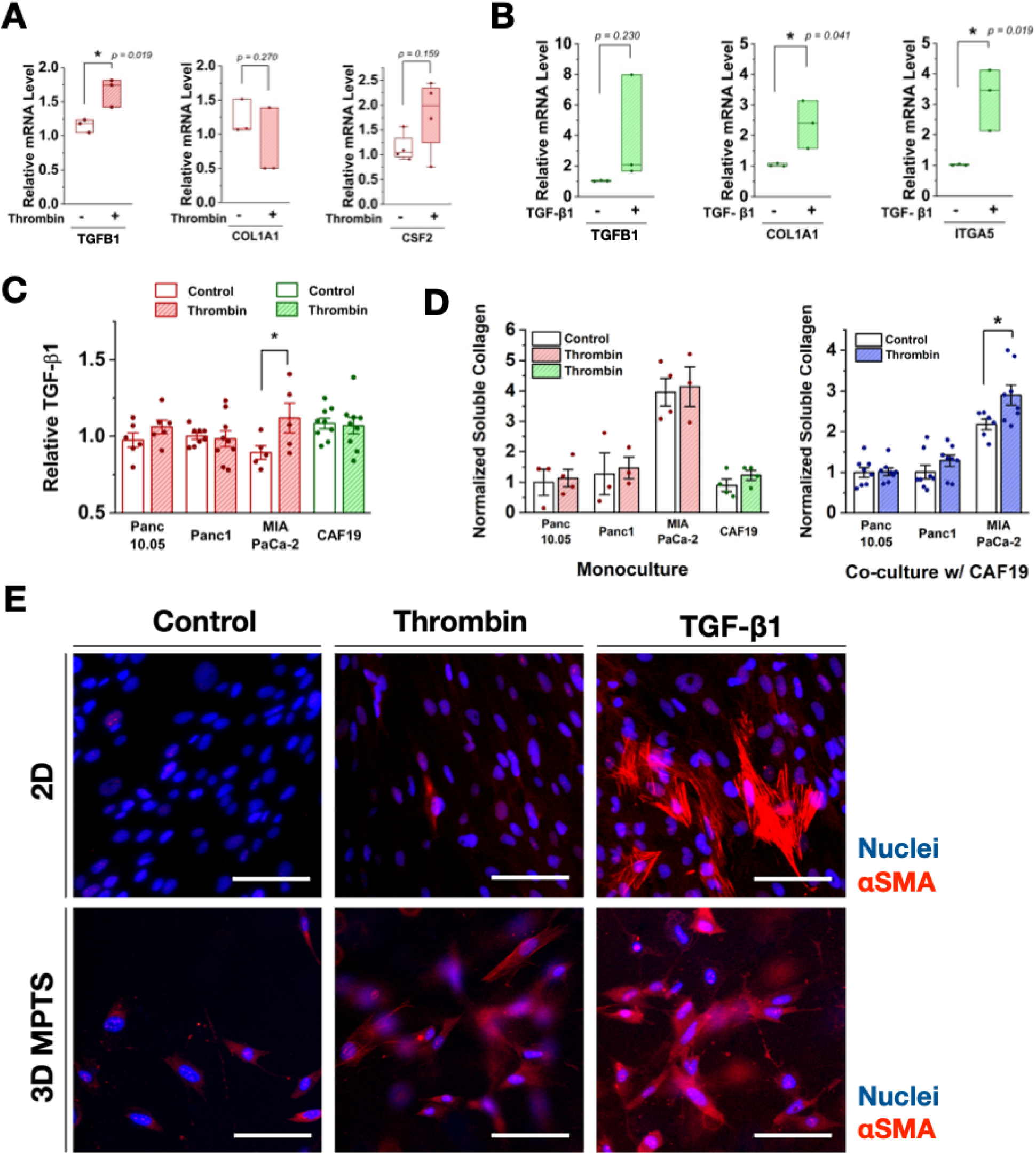
**Tumor-derived TGF-**β **signaling and stromal activation in human PDAC models.** (A) Quantitative PCR analysis of TGFB1, COL1A1, and CSF2 expression in MIA PaCa-2 cells with and without thrombin treatment. (B) Quantitative PCR analysis of fibrotic gene expression in CAF19 following thrombin or TGF-β1 treatment. (C) ELISA measurement of total TGF-β1 secretion from MIA PaCa-2 and CAF19 under thrombin stimulation. (D) ELISA quantification of soluble collagen levels in monoculture (left) and co-culture (right) conditions with and without thrombin treatment. (E) Immunofluorescence imaging of αSMA expression in CAF19 under thrombin or TGF-β1 treatment in 2D and 3D culture conditions. Scale bar = 100 μm. Statistical analysis with Student’s t test (N ≥ 3).

We next examined whether thrombin or TGF-β1 directly activates CAFs. In contrast to tumor cells, thrombin treatment did not significantly alter fibrotic gene expression in CAF19 (**Fig. S6**), separating stromal activation from direct thrombin signaling. Given the induction of TGFB1 in tumor cells and the minimal direct response of CAFs to thrombin, we tested whether tumor-derived TGF-β1 could mediate stromal activation. CAF19 cells were treated with exogenous TGF-β1 (**Fig. 5B**), which significantly increased expression of COL1A1 and ITGA5, consistent with enhanced fibrotic activity and integrin-mediated signaling [41, 42]. TGFB1 expression was also upregulated in CAFs following TGF-β1 stimulation, indicating a reinforcing signaling loop.

To further evaluate this mechanism, we measured secretion of total (latent and active) TGF-β1 by ELISA. Thrombin treatment increased TGF-β1 secretion in MIA PaCa-2 cells, whereas other PCCs and CAF19 showed minimal changes (**Fig. 5C**). This effect was again most pronounced in MIA PaCa-2, consistent with its high PAR1 expression, positioning tumor cells as the dominant source of TGF-β1 within this system. We next assessed downstream functional consequences by measuring extracellular matrix production. ELISA analysis of soluble collagen revealed that MIA PaCa-2-CAF19 co-cultures exhibited the highest collagen levels and the greatest increase following thrombin treatment, whereas monoculture conditions showed minimal changes (**Fig. 5D**), linking ECM remodeling to PAR1-dependent tumor–stroma coupling.

Immunofluorescence analysis further resolved CAF activation states. αSMA expression increased in CAF19 following direct thrombin exposure and TGF-β1 treatment (**Fig. 5E**). TGF-β1 induced a stronger increase than thrombin alone, placing tumor-derived TGF-β1 downstream of PAR1 as the dominant driver of CAF activation. This pattern was consistent across other CAF cell lines. Comparable responses were observed in a different human CAF line (02-hT), demonstrating that TGF-β1-mediated stromal activation is not restricted to a single CAF model (**Fig. S7**). The results suggest that thrombin-PAR1 signaling in tumor cells induces TGF-β1 production and secretion, which activates CAFs and drives extracellular matrix remodeling through a paracrine mechanism embedded within the tumor-stroma-coagulation axis.

### PAR1 inhibition suppresses tumor-stroma interactions and stromal growth across human PDAC models

To determine whether PAR1 signaling is required for maintenance of tumor-stroma coupling within the tumor-stroma-coagulation axis, we evaluated the effects of pharmacologic PAR1 inhibition using a PAR1 antagonist, vorapaxar (VPX). First, cytotoxicity of VPX was assessed in MIA PaCa-2 and CAF cells, confirming that changes in cell proliferation are attributable to PAR1 inhibition rather than nonspecific cytotoxic effects (**Fig. S8**).

Human MPTS of PDAC were treated with 0, 5, and 10 µM VPX and continuously exposed to thrombin across all treatment groups for eight days. Fluorescence micrographs showed a dose-dependent reduction in both PCC and CAF cell density across all PCC-CAF co-culture conditions (**Fig. 6A**). Quantitative analysis demonstrated that VPX treatment significantly suppressed CAF growth across all co-culture models (**Fig. 6B**). PDAC cell growth was also reduced, but to a lesser extent than CAFs. Among the tumor cell lines, MIA PaCa-2, which exhibits the highest PAR1 expression, showed a statistically significant reduction in growth, decreasing by approximately 0.7-fold at the highest VPX concentration, linking sensitivity to tumor-intrinsic PAR1 levels. The preferential suppression of CAF growth, together with attenuation of PCC expansion under co-culture conditions, positions stromal reprogramming, rather than direct suppression of tumor cell proliferation, as the dominant consequence of PAR1 inhibition within the tumor-stroma-coagulation axis.

**Figure 6.**
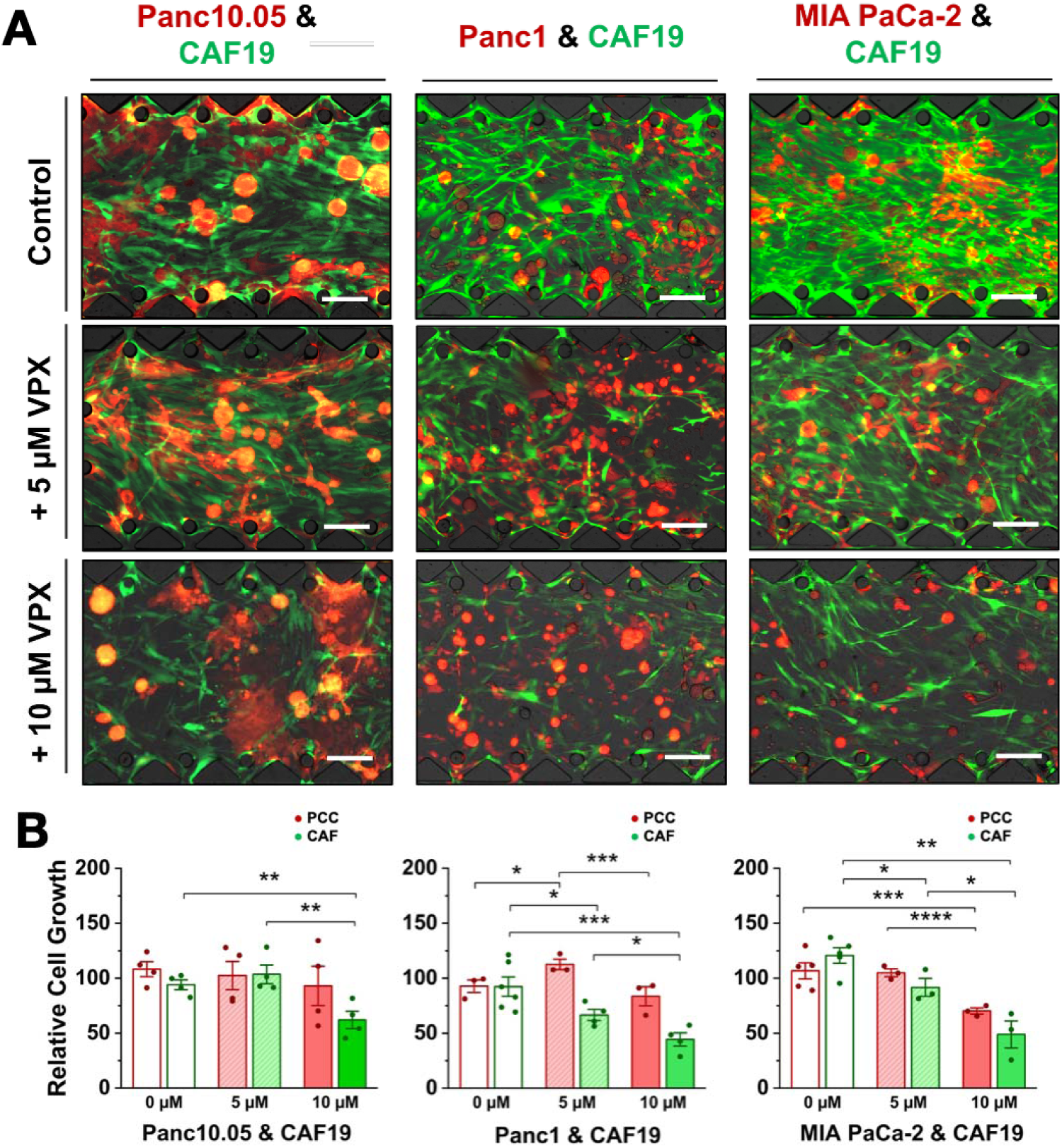
PAR1 inhibition suppresses PAR1-dependent tumor–stroma interactions in human PDAC co-culture models. (A) Representative fluorescence micrographs of PCC–CAF co-cultures (Panc10.05-CAF19, Panc1-CAF19, and MIA PaCa-2-CAF19) treated with 0, 5, and 10 µM vorapaxar (VPX) under continuous thrombin stimulation. Scale bar = 200 μm. (B) Quantification of cell growth in PCC-CAF co-culture conditions following VPX treatment at Day 8. Statistical analysis with one-way ANOVA with post hoc Tukey test (N ≥ 3).

### PAR1 inhibition reprograms fibrosis-associated stromal states in vivo

To determine whether the stromal reprogramming observed in vitro extends in vivo, we performed spatial transcriptomic analysis of orthotopic KPC tumors following daily treatment with VPX (15 mg/kg) for 14 days (**Fig. 7A**). Tumors were harvested at the endpoint for histological and spatial profiling. VPX treatment did not significantly alter tumor weight compared with control animals (**Fig. 7B**), contrasting with previously reported effects of tumor-intrinsic Par1 deletion [22]. This difference likely reflects mechanistic ablation from tumor initiation versus pharmacologic perturbation after establishment of the TME, when CAF populations, ECM deposition, and TGF-β1-dependent feedback may partially sustain stromal signaling. Thus, pharmacologic PAR1 inhibition preferentially reprograms established stromal states rather than producing immediate collapse of primary tumor burden.

**Figure 7.**
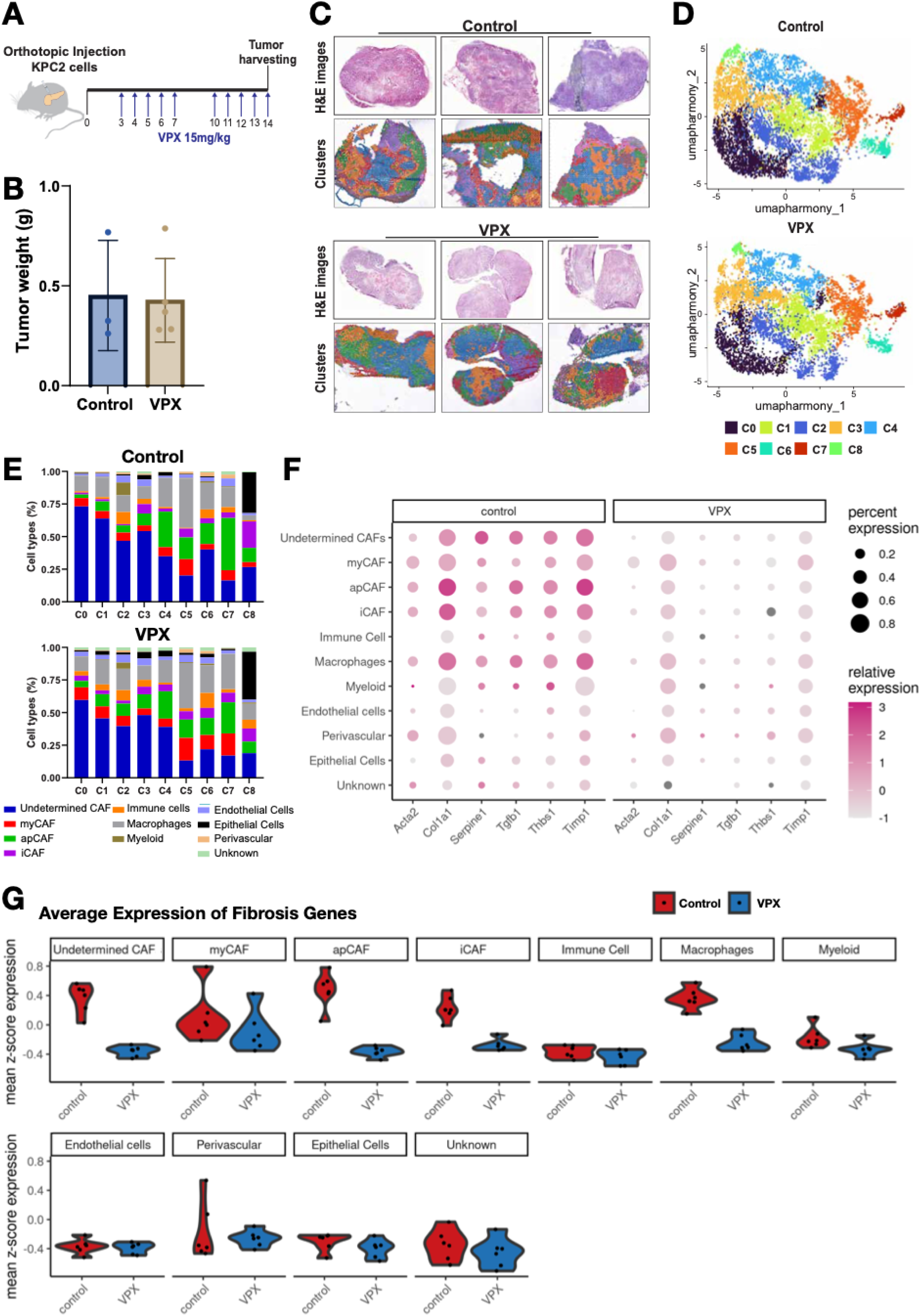
Spatial transcriptomic analysis of PAR1 inhibition in orthotopic PDAC tumors. (A) Experimental schematic showing orthotopic implantation of KPC tumors and daily VPX treatment (15 mg/kg) until tumor harvest at day 14. (B) Tumor weight in control and VPX-treated groups. (C) Representative H&E images and spatial clustering of control and VPX-treated tumors. (D) UMAP visualization of spatial transcriptomic clusters in control and VPX-treated tumors. (E) Cell-type annotation of spatial clusters, including CAF subpopulations. (F) Changes in fibrosis-associated gene expression following VPX treatments. (G) Average expression of fibrosis genes in each cell type.

Histological and spatial transcriptomic analyses revealed substantial reorganization of tumor architecture following VPX treatment. Spatial clustering analysis together with manual examination of H&E stained tissue sections (**Fig. S9**) demonstrated distinct spatial patterning of stromal tissues between control and VPX-treated tumors (**Fig. 7C**). Uniform manifold approximation and projection (UMAP) analysis showed reorganization of cell states in response to PAR1 inhibition (**Fig. 7D**), resolving a shift in tumor and stromal cell states rather than uniform suppression. Cell-type annotation of spatial clusters further defined these changes at the population level (**Fig. 7E**): VPX treatment reduced the relative proportion of multiple CAF populations, including general CAFs, inflammatory CAFs (iCAFs), and antigen presenting CAFs (apCAFs), indicating that PAR1 inhibition reshapes the composition of the stromal compartment rather than uniformly suppressing all CAF states. This restructuring of CAF proportions supports a central role for the tumor-stroma-coagulation axis in maintaining fibrosis-associated stromal architecture, and motivated a complementary analysis of whether these compositional shifts were accompanied by corresponding changes in fibrotic gene activity within each population.

To further resolve the impact of PAR1 inhibition on stromal transcriptional programs, we examined the expression of fibrosis-associated genes across spatially defined cell populations (**Fig. 7F**). VPX treatment reduced expression of key pro-fibrotic genes, including *Tgfb1*, *Acta2*, *Serpine1*, and *Thbs1*, most broadly across undetermined CAFs, iCAFs, and apCAFs, indicating that fibrotic gene attenuation is not restricted to a single transcriptionally defined subtype but extends across the majority of the annotated stromal compartment. Because undetermined CAFs constitute a substantial fraction of this responsive population, this broad attenuation should be interpreted primarily as a population-level suppression of fibrotic signaling rather than a subtype-restricted effect, and we cannot exclude that part of the undetermined-CAF response reflects cells transitioning between, or incompletely resolved among, defined subtypes. Against this broadly responsive background, myCAFs exhibited a distinct pattern: fibrotic gene expression was reduced within individual cells, but the proportion of myCAFs expressing these markers was unchanged, indicating that PAR1 inhibition suppresses ongoing transcriptional activity in established myCAF populations without altering myCAF abundance or spatial prevalence. This contrast indicates differential sensitivity to PAR1 inhibition across the CAF compartment, with most subtypes (and the undetermined population) responding through reduced representation or broad transcriptional attenuation, while myCAFs respond selectively at the level of fibrotic gene activity within a stable population. Average expression profiles across cell types further support this pattern of coordinated, but subtype-differentiated, suppression of fibrotic programs (**Fig. 7G**).

### PAR1 inhibition reprograms tumor-stroma interactions in human PCC-CAF xenograft models

To evaluate whether thrombin-PAR1-driven stromal activity within the tumor-stroma-coagulation axis is maintained in in vivo models with human tumors, we established a PCC-CAF xenograft model. Mice were subcutaneously injected with TdTomato-labeled MIA PaCa-2 cells and GFP-labeled CAF19 cells at a 1:5 ratio. This ratio was optimized based on endpoint analyses of CAF persistence, accounting for their lower proliferation and survival relative to tumor cells (**Fig. S10**). Immunofluorescence staining confirmed establishment of both cell types containing substantial populations of PCCs (RFP+) and CAFs (GFP+) after four weeks of growth (**Fig. 8A** and **8B**).

**Figure 8.**
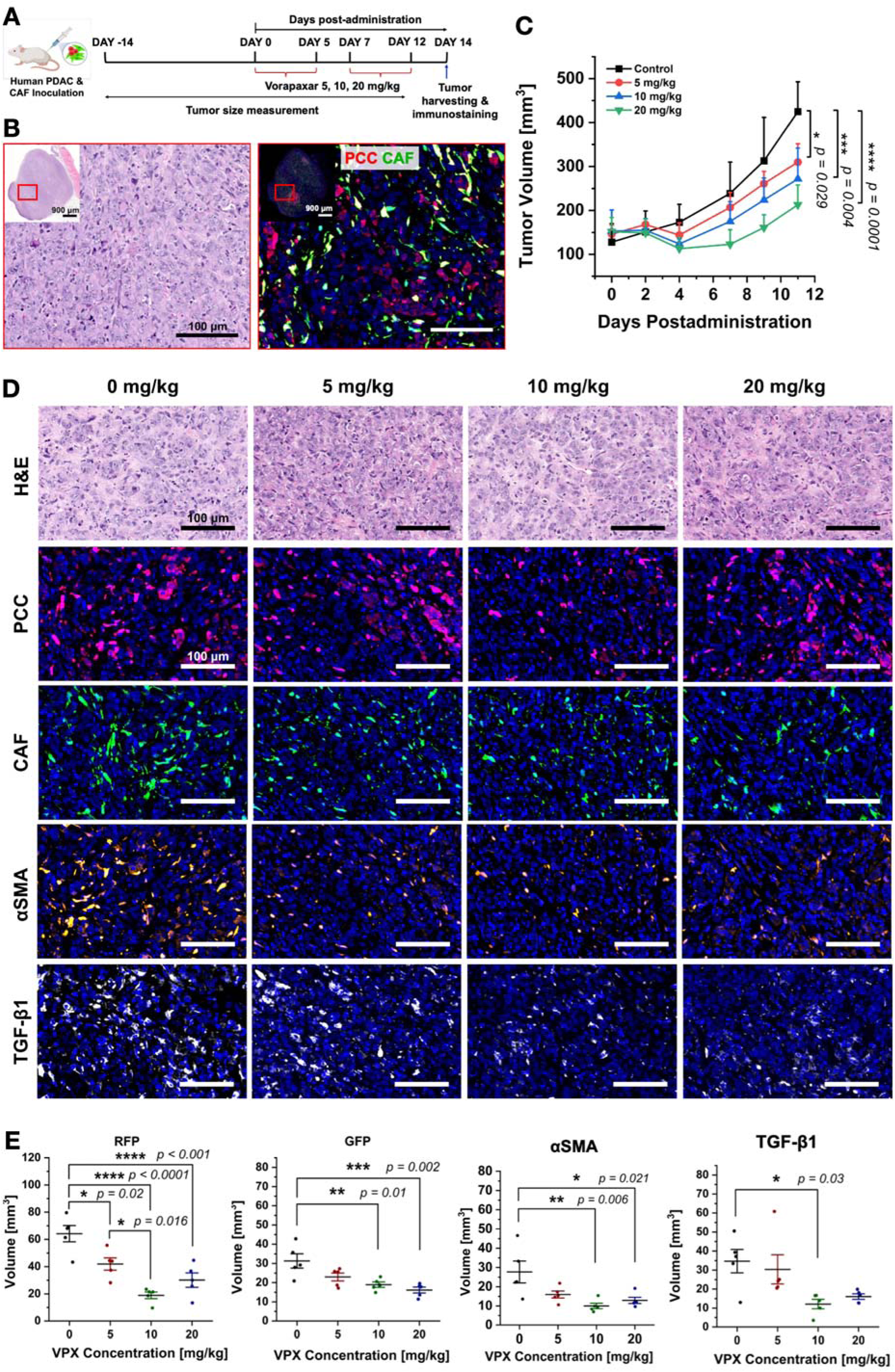
PAR1 inhibition in human PCC-CAF xenograft models. (A) Experimental schematic showing xenograft establishment using TdTomato-labeled MIA PaCa-2 cells and GFP-labeled CAF19 cells and VPX treatment schedule. (B) H&E and immunofluorescence staining of PCCs (red) and CAFs (green) in tumor sections. Scale bar = 100 µm. (C) Normalized tumor volume following VPX treatment. (D) Immunofluorescence staining of PCCs, CAFs, αSMA, and TGF-β1 in tumor sections. Scale bar = 100 µm. (E) Quantification of stained areas for PCCs, CAFs, αSMA, and TGF-β1. Statistical analysis with one-way ANOVA with post hoc Tukey test (N ≥ 3).

To allow tumor establishment prior to treatment, xenografts were grown for 14 days before initiation of VPX administration with an average tumor volume of 145 ± 35.5 mm^3^. VPX was delivered via oral gavage for an additional 14 days using two cycles of five days of treatment followed by two days of rest. VPX administration produced a dose-dependent reduction in tumor volume, with partial rebound following the initial treatment cycle (**Fig. 8C**), consistent with a reversible microenvironmental response rather than sustained cytotoxic tumor control.

Beyond tumor size, immunofluorescence analysis resolved compartment-specific effects. VPX treatment reduced both tumor (RFP+) and stromal (GFP+) compartments in a dose-dependent manner (**Fig. 8D** and **8E**). Quantitative analysis further showed decreased expression of αSMA and TGF-β1, aligning with suppression of CAF activation and attenuation of TGF-β–dependent signaling. This reduction in stromal activation is consistent with disruption of the PAR1–TGF-β signaling cascade defined in earlier experiments and reflects coordinated reprogramming of tumor and stromal compartments.

## DISCUSSION

PDAC is characterized by a highly desmoplastic tumor microenvironment in which stromal components exert both tumor-promoting and tumor-restraining functions [4, 43]. This duality has limited the success of strategies aimed at broadly depleting the stroma, highlighting the need for approaches that selectively reprogram stromal states. In this study, extravascular coagulation emerges as a regulatory module that stabilizes pro-fibrotic stromal states through tumor-intrinsic PAR1 signaling. Integration of human tumor data, microphysiological models, and in vivo validation defines a tumor-stroma-coagulation axis that links coagulation signaling to stromal remodeling and tumor progression.

Coagulation signaling is commonly interpreted as a consequence of tumor-associated vascular leakage [44, 45]. The present results instead position coagulation as an upstream regulator of tumor-stroma communication, extending a conceptual lineage first articulated in the "tumors as wounds that do not heal" framework [45]. That framework proposed that tumors co-opt wound-healing programs, including coagulation and fibrin deposition, to sustain a permissive stroma. Our findings extend this concept mechanistically: coagulation does not merely deposit fibrin as a passive scaffold but actively initiates downstream stromal remodeling through tumor-intrinsic PAR1 signaling, which in turn induces TGF-β1 to amplify stromal activation. Elevated F2R (PAR1) expression in human PDAC associates with fibrotic transcriptional programs and poor survival, while pathway-level analysis resolves coordinated activation of ECM, TGF-β, and coagulation signaling programs. These programs dominate over coagulation itself at the transcriptional level, placing coagulation as a regulatory input that organizes downstream stromal remodeling, rather than a parallel bystander process.

Mechanistically, PAR1 activation drives stromal remodeling through induction of tumor-derived TGF-β1. Thrombin-PAR1 signaling induces TGFB1 secretion from tumor cells, whereas CAFs show only weak direct responses to thrombin but activate robustly upon TGF-β1 exposure, driving ECM deposition and fibroblast activation. This establishes PAR1 as the control node initiating a directional, tumor-to-CAF cascade that is then amplified as activated CAFs induce their own TGFB1, creating a self-sustaining loop within the TGF-β arm that reinforces fibrotic stromal states downstream of the initiating PAR1 signal [46–48]. Additional components identified through transcriptomic analysis, including THBS1, FURIN, and SERPINE1, map onto processes of TGF-β1 activation [32, 34, 35], and fibrinolysis inhibition and subsequent matrix stabilization [37, 38]. These factors extend the signaling cascade into a broader regulatory network that sustains fibrosis. The resulting increase in activated TGF-β1 and expansion of the CAF population can collectively promote PDAC progression, persistent fibrosis-associated stromal remodeling and tumor-supportive microenvironmental states [49–52]. Within this framework, coagulation signaling initiates a tumor-stroma amplification circuit that is reinforced through multiple downstream modules and stabilizes a pro-fibrotic microenvironment. Notably, thrombin can modestly influence CAF behavior directly, but tumor-mediated activation via paracrine TGF-β1 signaling dominates, reinforcing the importance of targeting this upstream node rather than individual downstream effectors.

The spatial transcriptomic data further reveal that PAR1 inhibition engages the CAF compartment non-uniformly, consistent with established heterogeneity in CAF subtype biology [53, 54]. Most CAF subtypes, including undetermined CAFs, iCAFs, and apCAFs, responded to VPX treatment through reduced representation within the stromal compartment. In contrast, myCAFs exhibited a distinct pattern: fibrotic gene expression was reduced within individual cells without a corresponding change in myCAF abundance. This dissociation between transcriptional activity and population size suggests that PAR1 inhibition suppresses the fibrotic gene program actively maintained within established myCAFs, without eliminating the myCAF population itself. This pattern is consistent with the model in which myCAF and iCAF states are governed by distinct upstream signals including TGF-β and IL-1/JAK-STAT, respectively, which shape CAF identity in PDAC. Our data suggest that tumor-derived, PAR1-dependent TGF-β1 signaling specifically sustains the transcriptional output of the myCAF program, such that its pharmacologic disruption attenuates myCAF activity without depleting the population. This selective transcriptional reprogramming, rather than population-level elimination, is consistent with the goal articulated in the Introduction: disrupting tumor-promoting stromal function while preserving the stromal architecture itself.

Beyond defining the biological mechanism, this study illustrates how engineered microphysiological environments can resolve regulatory interactions that are difficult to isolate in conventional tumor models. The MPTS platform reconstructs tumor-stroma organization within a three-dimensional collagen matrix while providing controlled exposure to extravascular coagulation signals and compartment-resolved genetic and pharmacologic perturbation. This molecular and microenvironmental control enabled thrombin-PAR1 signaling to be separated from the broader complexity of the tumor microenvironment and revealed how tumor-intrinsic signaling is propagated into fibrosis-associated stromal responses. Importantly, comparison across murine and human MPTS models showed that conservation of the regulatory axis does not require identical phenotypic outputs: murine systems revealed strong PAR1-dependent tumor growth, whereas human models exposed context-dependent stromal activation gated by basal tumor PAR1 expression. Together with orthotopic and xenograft validation, these findings support engineered microphysiological systems as complementary models for resolving both mechanistic causality and biologically relevant heterogeneity across experimental scales.

The clinical associations observed in this study further support the relevance of the tumor-stroma-coagulation axis in PDAC progression. Elevated F2R expression correlates with reduced disease-specific survival, with divergence of survival curves occurring within the 12∼30 month interval, consistent with patterns of early recurrence reported following pancreaticoduodenectomy. Prior clinical studies [55, 56] have shown that perioperative therapy timing critically influences outcomes within this window, suggesting that early disease progression is driven by biologically aggressive tumor states. The association between high F2R expression and adverse outcomes provides a molecular framework for these observations, linking coagulation signaling to tumor-promoting stromal architecture. That survival tracks with F2R status rather than TGFB1 co-expression (**Fig. 1F**) reinforces PAR1 as the upstream determinant of clinical outcome, consistent with its role as the control node of this axis, and suggests that F2R expression alone, rather than a combined F2R/TGFB1 signature, may be the more appropriate variable for risk stratification and selection of patients for PAR1-targeted intervention.

Pharmacologic inhibition of PAR1 disrupts this circuit at its control node. In this study, vorapaxar served as a selective pharmacological probe establishing that PAR1 inhibition is sufficient to reprogram fibrosis-associated stromal states: VPX suppresses CAF proliferation, reduces αSMA expression, and attenuates ECM remodeling in vitro without inducing tumor cell cytotoxicity. In orthotopic KPC models, PAR1 inhibition reshapes the composition and transcriptional activity of the stromal compartment, with reduced proportions of several CAF subtypes and reduced fibrotic gene expression within myCAFs, while leaving overall tumor size largely unchanged. In the human PCC-CAF xenograft model, VPX reduces both tumor burden and stromal activation markers, confirming that disruption of the tumor-stroma-coagulation axis alters tumor progression across systems. Targeting PAR1 rather than TGF-β1 itself offers a practical advantage for therapeutic translation. Direct TGF-β pathway inhibition has already been tested clinically in PDAC, with galunisertib producing only modest survival benefit in combination with gemcitabine [57]. This limited clinical activity likely reflects TGF-β’s broader, context-dependent roles across normal and malignant tissue, which complicate systemic inhibition. PAR1 activation within the coagulation-active tumor interstitium may provide a more context-restricted entry point into the same downstream fibrotic program, while additionally engaging PAR1’s established roles in tumor cell proliferation and immune evasion, extending its therapeutic scope beyond stromal reprogramming alone.

Clinical translation of this mechanistic proof-of-concept requires distinguishing VPX’s role as a pharmacologic probe from its suitability as an oncologic therapeutic. VPX has a prolonged terminal half-life and near-irreversible PAR1 binding, resulting in sustained platelet inhibition that may limit dosing flexibility in patients with PDAC [58]. Notably, the partial rebound in tumor volume during the 2-day treatment-free intervals in the xenograft model (**Fig. 8C**) suggests that sustained PAR1 inhibition may be required to maintain stromal reprogramming. These findings motivate development of PAR1 inhibitors with pharmacokinetic properties better suited for oncology and evaluation of combination strategies that may enhance or prolong stromal reprogramming. The established feasibility of anticoagulation in patients with PDAC receiving chemotherapy further supports investigation of coagulation-axis-targeted therapeutic strategies [59].

While our study highlights thrombin-PAR1 signaling, it is important to recognize that PAR1 may be activated by multiple proteases. In addition to thrombin, other enzymes such as matrix metalloproteinases (MMPs) and plasmin have been reported to cleave PARs, raising the possibility that diverse proteolytic inputs converge on PAR-dependent signaling [60]. The impact of such alternative cleavage events on PCC-CAF crosstalk remains unexplored and warrants further investigation. Moreover, PAR1 is only one member of the PAR family and thrombin can cleave several PAR isoforms [60, 61], suggesting that overlapping or compensatory signaling may contribute to tumor-stroma interactions. Additionally, other cells in the PDAC stroma may also express PARs, including CAFs, platelets, immune, and endothelial cells, all of which could play critical roles in reprogramming the PDAC stroma and warrants further investigation [21–23]. These findings highlight the complexity of PAR biology in PDAC and point to the need for broader investigation into protease diversity and PAR family cross-regulation in mediating fibrosis.

Extravascular coagulation thus functions as a systems-level regulator of stromal state architecture in PDAC. PAR1 operates as a central control node that links coagulation signaling to tumor–stroma coupling and fibrosis-associated progression states. Targeting such regulatory nodes offers a strategy to selectively reprogram the PDAC stroma while preserving its tumor-restraining functions, providing a framework for therapeutic intervention grounded in microenvironmental regulation rather than tumor cell-intrinsic cytotoxicity.

## ACKNOWLEDGEMENT

The authors acknowledge Dr. Mark Band and Tatsiana Akraiko at the Functional Genomics Core of the Roy J. Carver Biotechnology Center. In addition, authors acknowledge Megan Cohen at the Purdue University Histology Research Laboratory for their histology, analysis, and tissue imaging. Authors also acknowledge Ffion Marie Edwards for VPX cytotoxicity assay. This work was partially supported by grants from the National Institutes of Health (U01 CA274304 to MJF/MLF/BH and R01 CA254110 to MLF/BH). We acknowledge the support from the Cancer Center at Illinois (P30 CA275774), Purdue Institute for Cancer Research (P30 CA023168), IU Simon Cancer Center (P30 CA082709), and the Walther Cancer Foundation. BH is a Biohub Investigator.

## DATA AVAILABILITY

All other raw data generated in this study are available upon request to the corresponding author.

## Notes

### Competing Interest Statement

The authors have declared no competing interest.

### Summary of Updates

Discussion and Figures have been updated.

