## Supplementary S1 for "Extravascular coagulation stabilizes pro-fibrotic stromal states via tumor-intrinsic PAR1 signaling in pancreatic ductal adenocarcinoma"

**SUPPLEMENTARY METHODS**

### RT-qPCR

A real-time quantitative polymerase chain reaction (RT qPCR)

To confirm the relative gene expression levels of Tissue Factor (TF) and PAR1 (F2R) for both murine and human cells lines, we performed RT qPCR of murine cell lines of KPC and mCAF, and human cell lines of Human Pancreatic Ductal Epithelial cells (HPDE), Panc10.05, Panc1, MIA PaCa-2, and CAF19. The HPDE cells were used as a control.

High quality RNA was isolated from each cell line using Direct-zol RNA Miniprep (Zymo Research CA, Catalog # R2051), following the manufacturer's guidelines. RNA concentration was determined using a NanoDrop™ One/OneC Microvolume UV-Vis Spectrophotometer (Thermo Scientific). 200 ng of total RNA was retrotranscribed and then amplified. RT-qPCR assays were performed with at least three biological replicates per gene, and each sample was analyzed in a minimum of two technical replicates using the PrimeTime One-Step RT-qPCR Master Mix (Integrated DNA Technologies, Catalog # 10007065) and Eukaryotic 18S rRNA as Endogenous Control (VIC™/MGB probe, Applied Biosystems, Catalog # 4319413E). The qPCR assays were run in the QuantStudio 7 Real-Time PCR System (Applied Biosystems). The data were analyzed and expressed relatively to the 18S rRNA in the thrombin stimulated sample and compared with the non-stimulated sample using the ΔΔCT method. Primer sequences are listed below:

Probes and Primer Sequences *Mus musculus* (Mm)*.

| **Gene Name** | **Oligonucleotide Type** | **Sequence** | **Ref Seq Number** |
| --- | --- | --- | --- |
| *Par1 F2r* | Probe | 5'/56-FAM/CTGTCTTCC/ZEN/CGCGTCCCTATGAG/3IABkFQ/-3' | NM_010169 Mm |
|  | Primer 1 | 5'-GGCGCTTGCTGATCGTC-3' |  |
|  | Primer 2 | 5'-CGTAGCATCTGTCCTCTCTGA-3' |  |
| *Lepr* | Probe | 5'-/56-FAM/TGAGGTATC/ZEN/ACAGGCGCAGCC/3IABkFQ/-3' | NM_146146 Mm |
|  | Primer 1 | 5'-TCACCCAGCACAATCCAAT-3' |  |
|  | Primer 2 | 5'-GCTCAGACGTAGGATGAATAGATG-3' |  |
| *Csf2* | Probe | 5'-/56-FAM/CGAATATCT/ZEN/TCAGGCGGGTCTGCA/3IABkFQ/-3' | NM_009969 Mm |
|  | Primer 1 | 5'-GTCTCTAACGAGTTCTCCTTCA-3' |  |
|  | Primer 2 | 5'-CCTTGAGTTTGGTGAAATTGCC-3' |  |
| *Cd44* | Probe | 5'-/56-FAM/TTCTTTATC/ZEN/CGGAGCACCTTGGCC/3IABkFQ/-3’ | NM_001177787  Mm |
|  | Primer 1 | 5'-ACCTTCCTACTGAACAGCCTA-3’ |  |
|  | Primer 2 | 5'-TGGAGTCCTTGGATGAGTCT-3’ |  |
| *Tgfb1* | Probe | 5'-/56-FAM/ATAGATGGC/ZEN/GTTGTTGCGGTCCA/3IABkFQ/-3' | NM_011577 Mm |
|  | Primer 1 | 5'-GCGGACTACTATGCTAAAGAGG-3' |  |
|  | Primer 2 | 5'-CCGAATGTCTGACGTATTGAAGA-3' |  |
| *Col1a1* | Probe | 5'-/56-FAM/CCGGAGGTC/ZEN/CACAAAGCTGAACA/3IABkFQ/-3' | NM-007742  Mm |
|  | Primer 1 | 5'-CGCAAAGAGTCTACATGTCTAGG-3' |  |
|  | Primer 2 | 5'-CATTGTGTATGCAGCTGACTTC-3' |  |

*Synthesized by Integrated DNA Technologies

Probes and Primer Sequences *Homo sapiens* (Hs)*.

| **Gene Name** | **Oligonucleotide Type** | **Sequence** | **Ref Seq Number** |
| --- | --- | --- | --- |
| *PAR1 F2R* | Probe | 5'/56-FAM/TGTATCCCA/ZEN/TGCAGTCCCTCTCCT/3IABkFQ/-3' | NM_001992 Hs |
|  | Primer 1 | 5'-CGCCTCTATCTTGCTCATGAC-3' |  |
|  | Primer 2 | 5'-GGCCAGACAAGTGAAGGAA-3' |  |
| *LEPR* | Probe | 5'/56-FAM/TGTTCCGAA/ZEN/CCCCAAGAATTGTTCCT/3IABkFQ/-3' | NM_002303  Hs |
|  | Primer 1 | 5'-TCACACCAAAGAATGAAAAAGCTA-3' |  |
|  | Primer 2 | 5'-CATGTCACTGATGCTGTATGC-3' |  |
| *CSF2* | Probe | 5'/56-FAM/CCAGCCACT/ZEN/ACAAGCAGCACTG/3IABkFQ/-3' | NM_000758  Hs |
|  | Primer 1 | 5'-CAGCCTCACCAAGCTCAAG-3' |  |
|  | Primer 2 | 5'-TGACAAGCAGAAAGTCCTTCAG-3' |  |
| *CD44* | Probe | 5'/56-FAM/ /3IABkFQ/-3' |  |
|  | Primer 1 | 5'- -3' |  |
|  | Primer 2 | 5'- -3' |  |
| *TGFB1* | Probe | 5'/56-FAM/ACCCGCGTG/ZEN/CTAATGGTGGAA /3IABkFQ/-3' | NM_000660  Hs |
|  | Primer 1 | 5'-CCGACTACTACGCCAAGGA -3' |  |
|  | Primer 2 | 5'-GTTCAGGTACCGCTTCTCG -3' |  |
| *COL1A1* | Probe | 5'/56-FAM/TCGAGGGCC/ZEN/AAGACGAAGACATC/3IABkFQ/-3' | NM_000088  Hs |
|  | Primer 1 | 5'-GACATGTTCAGCTTTGTGGAC-3' |  |
|  | Primer 2 | 5'-TTCTGTACGCAGGTGATTGG-3' |  |

*Synthesized by Integrated DNA Technologies

### RNA-seq and transcriptomic analysis

#### Cell preparation and RNA isolation

KPC tumor cells and mCAFs were processed under defined stimulation conditions. Cells were seeded at 1 × 10⁶ cells per 10 cm dish (day 0), followed by serum starvation in low-serum media (1% FBS) for 24 hours. Cells were then stimulated with thrombin (1 U/mL) or maintained as unstimulated controls for 24 hours (n = 3 per group). Cells were harvested in RNA preservation buffer (RNAprotect) to maintain RNA integrity.

#### Library preparation and sequencing

RNA samples were submitted to sequencing facilities (e.g., Purdue Genomics Core for KPC samples and Carver Functional Genomic Core for mCAF) for quality assessment and library preparation. Libraries were generated using standard poly(A)-enriched protocols and sequenced on Illumina platforms using paired-end reads (100–150 bp) with a minimum depth of 30 million reads per sample.

#### Quality control and preprocessing

For KPC samples, raw reads were processed using FastX-Toolkit (v0.0.13) with adapter trimming, minimum Phred score ≥30, and minimum read length ≥30 bp. For mCAF samples, preprocessing was performed using fastp (v0.23.2) with adapter trimming, minimum Phred score ≥30, and minimum read length ≥50 bp.

#### Read alignment and quantification

Quality-filtered reads were aligned to the mouse reference genome (GRCm38) using Tophat2 (v2.0.7) for KPC datasets and STAR aligner (v2.7.10a) for mCAF datasets. Gene-level counts were generated using HTSeq (v0.6.1) for KPC samples and FeatureCounts (Subread v1.6.1) for mCAF samples.

#### Differential expression analysis

Differential expression (DE) analysis was performed using the edgeR package (Bioconductor). For KPC datasets, genes with FDR ≤ 0.01 and absolute fold-change ≥2 were considered significant. For mCAF datasets, differential expression was defined using FDR < 0.05 following Benjamini–Hochberg correction.

#### Pathway and gene set enrichment analysis

Significant genes were analyzed using Ingenuity Pathway Analysis (IPA; QIAGEN) to identify enriched signaling pathways. Gene Set Enrichment Analysis (GSEA) was performed using the Broad Institute GSEA tool with MSigDB gene sets to assess enrichment of pathways associated with coagulation, TGF-β signaling, extracellular matrix organization, and related processes.

#### Clustering and visualization

Co-expressed gene clusters were identified using Clust-based approaches. Heatmaps were generated using the R package pheatmap to visualize differential gene expression across experimental conditions.

### MPTS device fabrication and operation

PDMS devices were fabricated using soft lithography. PDMS (10:1 base:curing agent) was cast on silicon molds and cured at 80°C for 4 h. Devices were bonded to glass slides after oxygen plasma treatment and baked at 120°C. Channels consisted of 300 μm source/sink channels connected by a 1 mm interstitial channel (100 μm height). Devices were sterilized using UV/ozone. Type I collagen (6 mg/mL) was prepared using PBS, NaOH, HEPES, FBS, and supplements. Tumor or tumor–CAF mixtures (1:1) were embedded at 2 × 10⁶ cells/mL. Collagen was polymerized at 37°C for 1 h. Thrombin (0.2 U/mL) and vorapaxar (5–10 μM) were introduced via perfusion with hydrostatic pressure differences. Cultures were maintained for 8 days.

### Cell growth quantification

Fluorescence images were acquired at multiple time points. Cell area was quantified using Trainable Weka Segmentation in FIJI ImageJ as illustrated next page. Growth was normalized to control conditions as below.

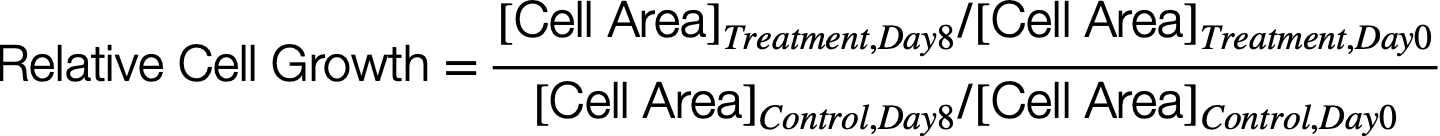

### TGF-β1 ELISA

Cells were seeded in 96-well plates (2000 cells/well; co-culture 1000:1000). After 5 days with or without thrombin, supernatants were collected and analyzed using Human TGF-β1 DuoSet ELISA (R&D Systems).

### Collagen secretion assay

Cells were seeded in 24-well plates (4000 cells/well). After 5 days, supernatants were analyzed using the Sircol soluble collagen assay (Biocolor).

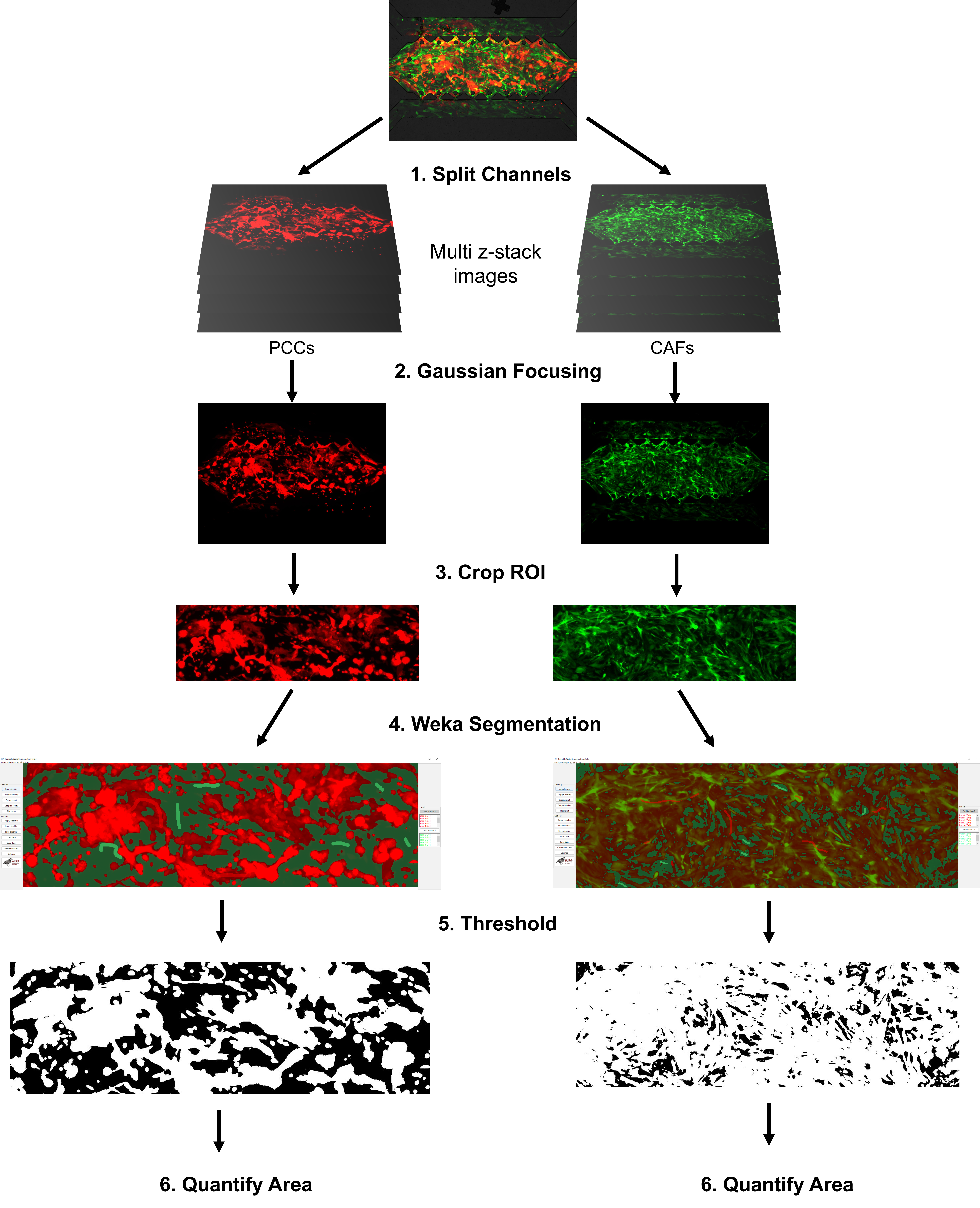

**Supplementary Method Figure 1.** Image processing method in FIJI ImageJ using Trainable Weka Segmentation method for cell area quantification.

### Spatial transcriptomics and treatment

#### Animal model and orthotopic tumor implantation

All animal experiments were conducted in accordance with institutional guidelines and approved protocols by Purdue University. C57BL/6 mice were used for orthotopic implantation of KPC tumor cells and were randomly assigned to vehicle control (n = 3) or vorapaxar (VPX) treatment groups (n = 5). KPC2 cells were resuspended at a density of 100,000 cells/mL in sterile PBS and implanted orthotopically into the pancreas under anesthesia using standard surgical procedures. At endpoint, tumor weights were measured. Hepatic metastatic foci were enumerated by gross inspection of the liver surface.

#### Vorapaxar (VPX) treatment

VPX was administered by oral gavage at a dose of 15 mg/kg. Treatment was initiated 3 days following tumor implantation and continued daily until tumor harvest.

#### Tumor harvest and tissue processing

At the experimental endpoint, mice were euthanized and tumors were surgically excised. Tissues were flash-frozen in isopentane (2-methylbutane) pre-cooled in liquid nitrogen and stored at −80 °C. Cryosections were prepared and mounted onto Visium spatial gene expression slides.

#### Spatial transcriptomics library preparation and sequencing

Spatial gene expression profiling was performed using the 10x Genomics Visium platform. Cryosections from flash-frozen tumor tissues were placed on Visium capture slides, with up to four sections per slide to enable parallel processing of biological replicates and conditions. Tissue sections were processed following the manufacturer’s protocol for fresh-frozen samples, including fixation, staining, permeabilization, reverse transcription, second-strand synthesis, cDNA amplification, and library preparation. Sequencing was performed on an Illumina platform to generate spatially barcoded transcriptomic data.

#### Image alignment and data processing

Histological images were aligned to capture areas using the Visium Image Alignment function in Loupe Browser (10x Genomics). Manual alignment was performed when necessary to correct for tissue positioning artifacts and ensure accurate correspondence between spatial barcodes and tissue morphology. Sequencing data were processed using the Space Ranger pipeline (10x Genomics), incorporating aligned image data to generate gene expression matrices linked to spatial coordinates.

#### Spatial transcriptomics data generation and preprocessing

Spatial transcriptomics data were generated using the 10x Genomics Visium platform and processed with the Space Ranger pipeline (version 2.1.1, 10X Genomics) to perform image alignment, barcode assignment, and gene expression quantification. Raw sequencing reads were aligned to the mouse (mm10) reference genome, and gene expression matrices for each spatial capture area were generated using default parameters.

#### Data import and quality control

Filtered gene-barcode matrices were imported into R and analyzed using Seurat ^1, 2^. Spot-level quality control was performed by examining standard metrics including total UMI counts, number of detected genes, percentage of mitochondrial gene expression, ribosomal gene content, and hemoglobin gene expression. Spots with extreme gene counts or high mitochondrial, ribosomal or hemoglobin content were excluded from downstream analyses based on dataset-specific thresholds.

#### Normalization and data integration

Data were normalized using Seurat’s SCTransform workflow ^3^. To account for batch effects across samples or experimental conditions, datasets were integrated using the Harmony algorithm ^4^ through Seurat’s layer-based integration framework. Harmony integration was performed on principal component embeddings derived from SCTransform-normalized data, generating corrected low-dimensional representations for downstream analyses.

#### Dimensionality reduction and clustering

Cell–cell neighborhood graphs were constructed using Harmony-corrected embeddings, followed by graph-based clustering implemented in Seurat. Low-dimensional visualization was performed using Uniform Manifold Approximation and Projection (UMAP) based on the integrated embeddings, with resolution parameters optimized to capture biologically meaningful clusters.

#### Cell-type annotation and spatial analysis

Cell type annotation was performed using a combination of automated and manual approaches. Initial annotations were assisted using SingleR, which provided reference-based predictions of cell identities. Final cell type annotations were assigned in a semi-automated fashion by integrating SingleR predictions ^5^ with cluster-specific marker gene expression and established canonical markers ^6^ , ensuring biologically meaningful classification of cell populations.

Cell-type annotation was performed using reference transcriptomic datasets, including MouseRNAseqData and Immunological Genome Project (ImmGen) panels, combined with manual curation based on established marker genes from published single-cell RNA-seq and spatial transcriptomics studies.

Analysis focused on three major compartments:

- Stromal cells, including undetermined CAFs, inflammatory CAFs (iCAFs), antigen-presenting CAFs (apCAFs), and myofibroblastic CAFs (myCAFs)
- Tumor cells, including epithelial, acinar, and ductal populations
- Immune cells, including dendritic cells, monocytes, neutrophils, T cells, NK cells, and myeloid populations

Macrophages and myeloid cells were analyzed as distinct populations due to their functional relevance. Annotation and visualization were performed using Loupe Browser, supported by marker gene expression patterns and spatial localization.

#### Differential expression analysis

Differential expression analysis was performed across spatial clusters and treatment conditions to identify genes modulated by VPX treatment. Gene expression changes were analyzed within defined cell populations and spatial regions to resolve treatment-associated transcriptional shifts.

Gene expression patterns were visualized using Seurat and dittoSeq ^7^. Tretment-level expression patterns were summarized using dot plots for specific marker genes. Signature (coagulation, fibrosis and immune suppression) expression across annotated cell types was visualized using the dittoSeq function dittoPlotVarsAcrossGroups, which computes average expression of given genes as a signature and summarizes across groups and enables stratified comparisons by treatment. Custom visualizations, including faceted representations, custom colors and related customizations were assisted by ggplot2-based approaches.

#### Statistical analysis

Statistical comparisons between experimental groups and identification of differentially expressed genes were performed using Seurat’s differential expression framework. By default, Seurat applies a two-sided non-parametric Wilcoxon rank-sum test to compare gene expression between groups of cells. Adjusted p-values were calculated using Bonferroni correction across all tested genes, as implemented in Seurat. Genes with adjusted p-values < 0.05 were considered statistically significant where specified.

#### Software and tools

All R-based were performed in R (version 4.3.3) using Seurat (version 5.1.0), Harmony (version 1.2.0), singleR (version 2.4.1), celldx (version 1.12.0), clustree (version 0.5.1) ^8^, dittoSeq (version 1.14.3), loupeR (version 1.1.0), tidyverse (version 2.0.0) and ggplot2 (version 3.5.1) packages.

**SUPPLEMENTARY FIGURES**

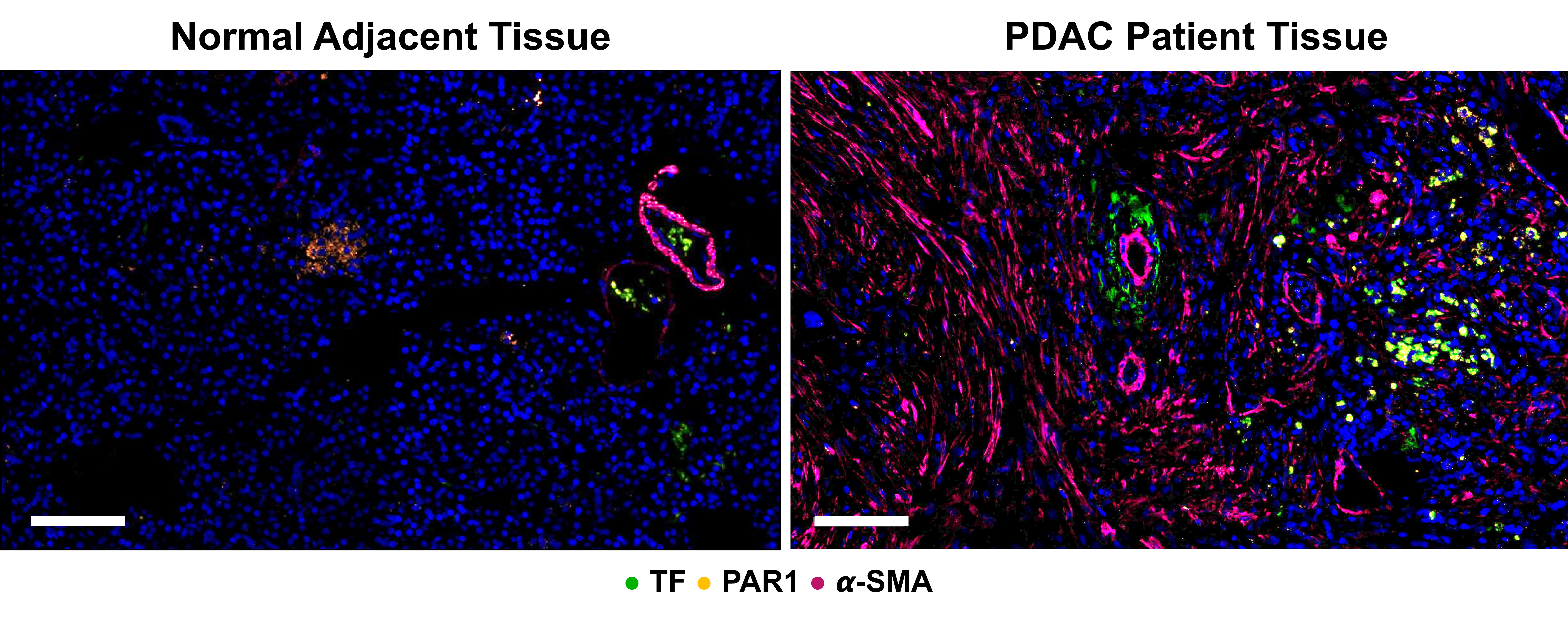

**Individual Fluorescence Channels of PDAC Patient Tissue #1**

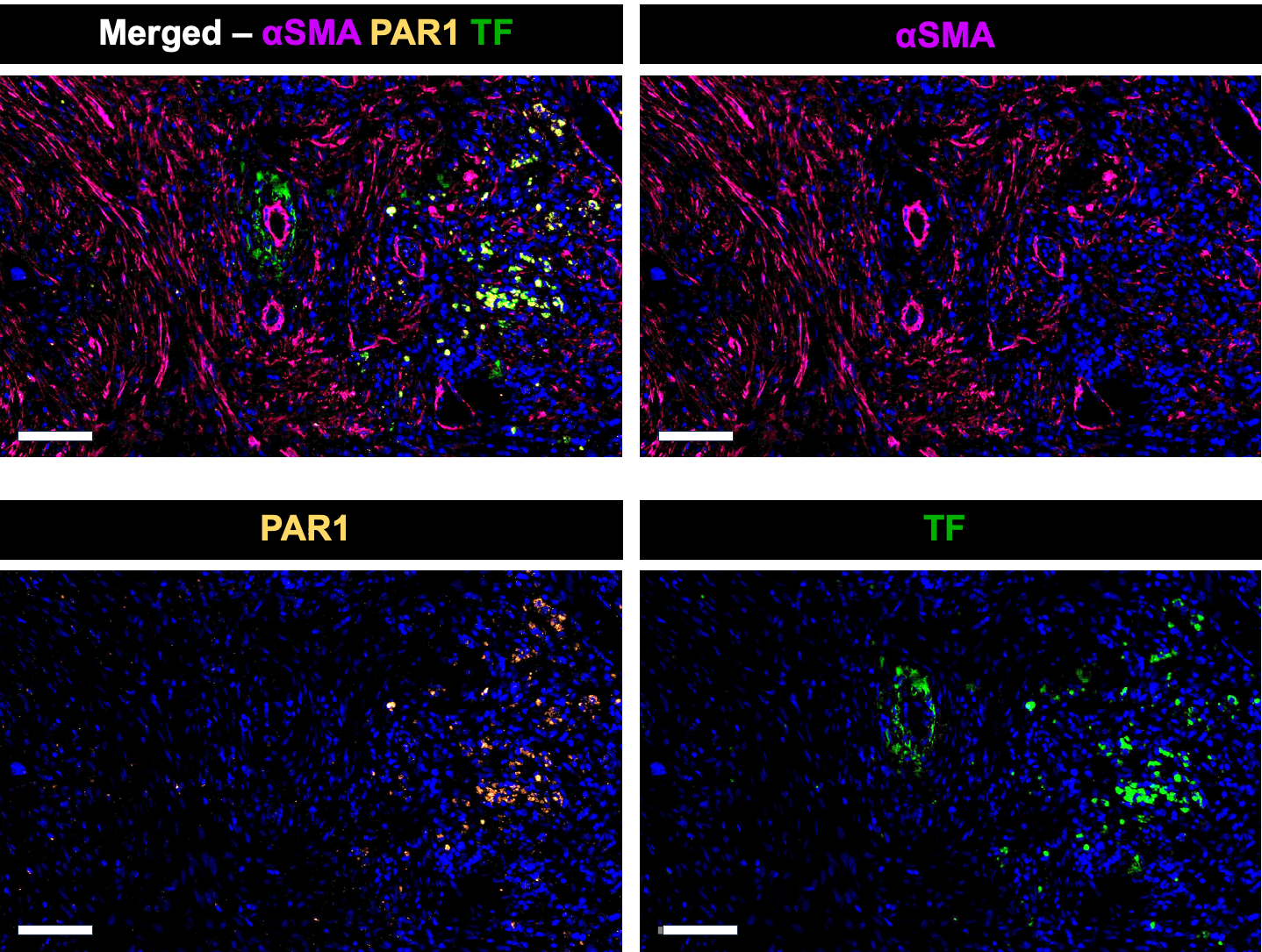

Scale bar = 100 μm

**Individual Fluorescence Channels of PDAC Patient Tissue #2**

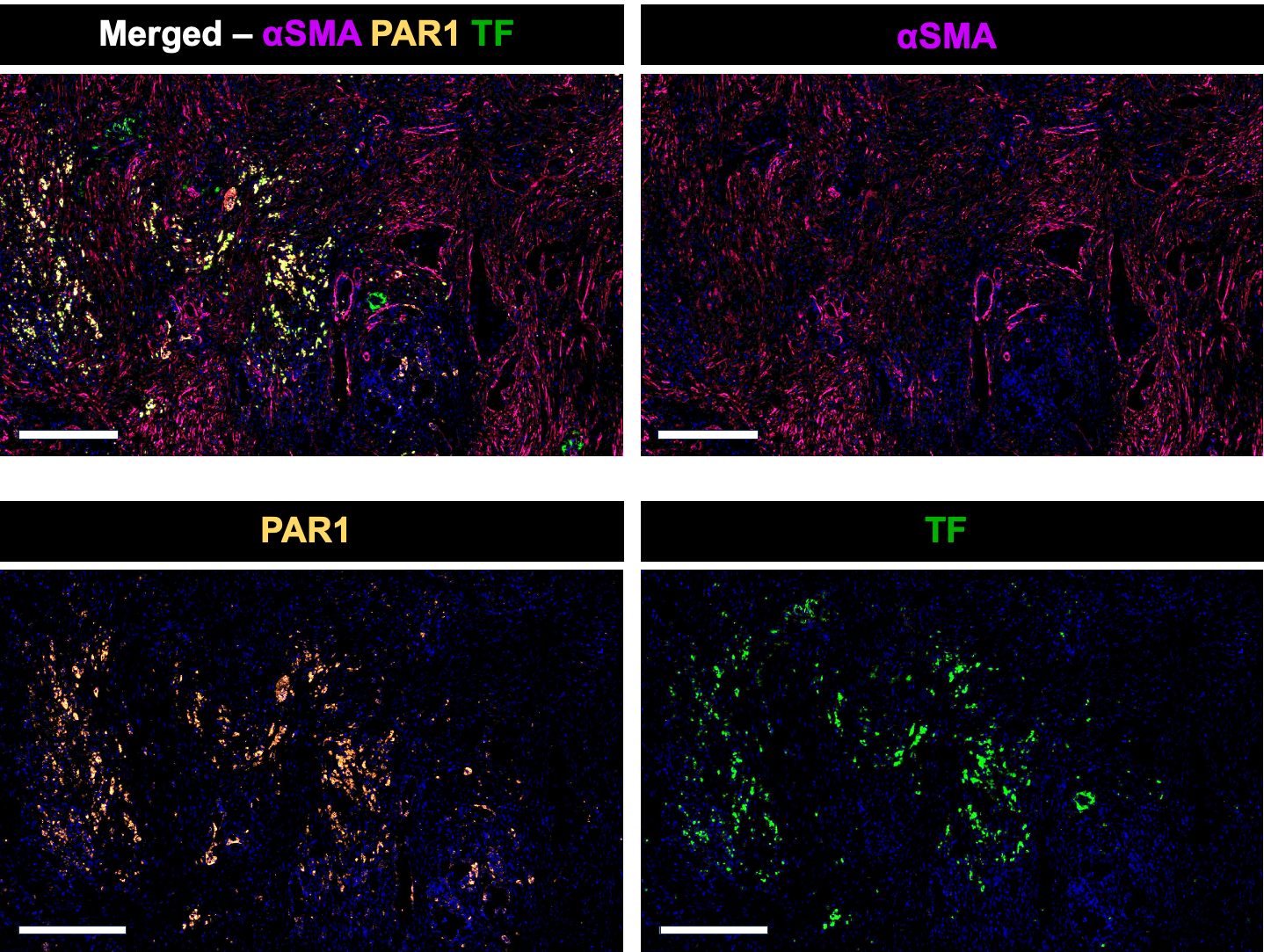

Scale bar = 300 μm

**Figure S1. Coagulation factor expressions** **human pancreas tissue samples** in normal pancreata and PDAC tissue from patient samples.

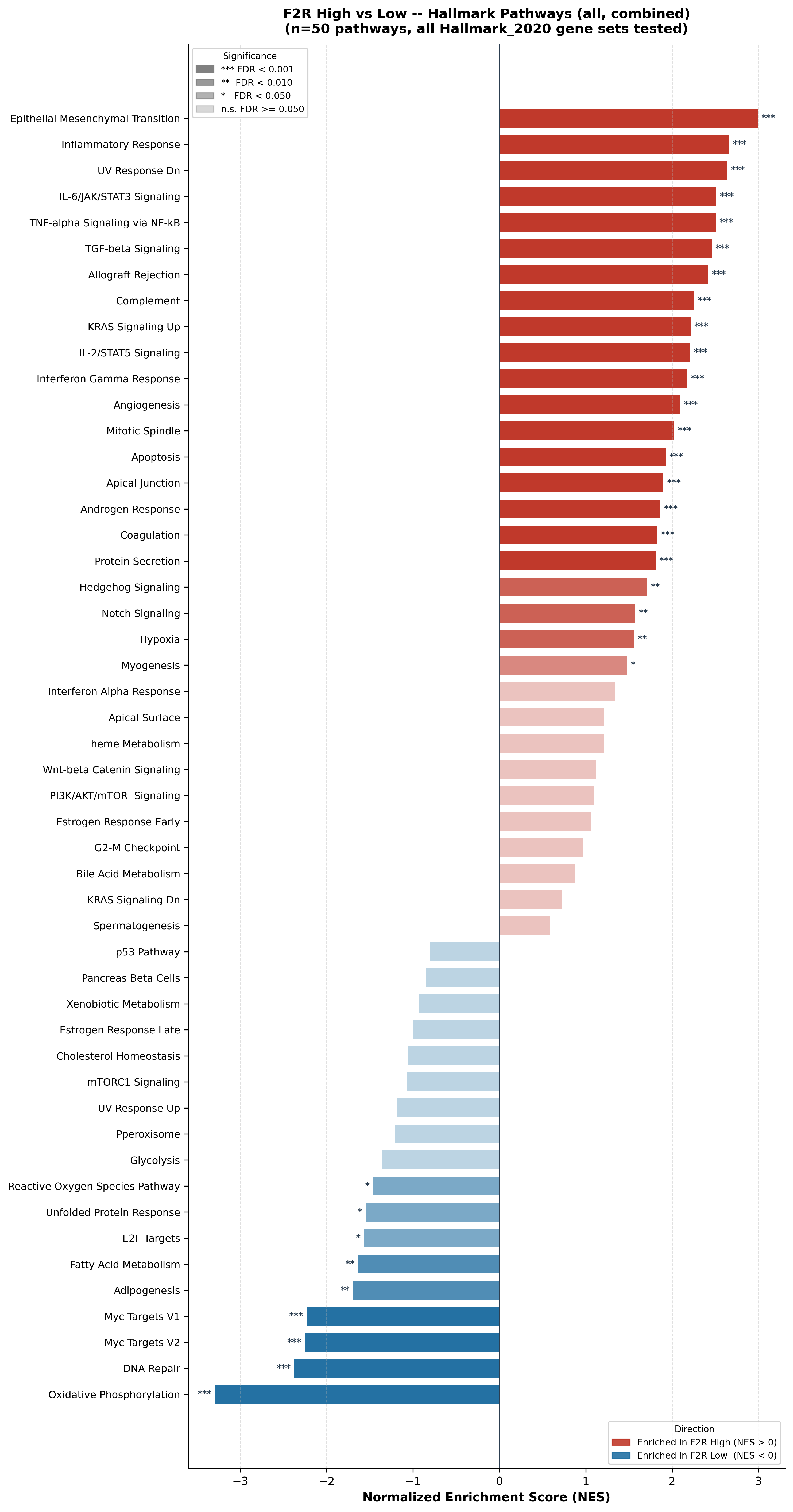

**Figure S2.** GSEA of F2R-High vs F2R-Low TCGA cohort by the Hallmark gene set.

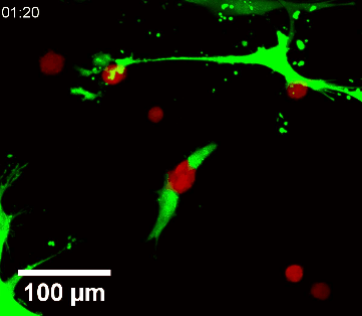

**Supplementary Video 1. Time-lapse micrograph of PCC-CAF interaction on the T-MOC platform**.

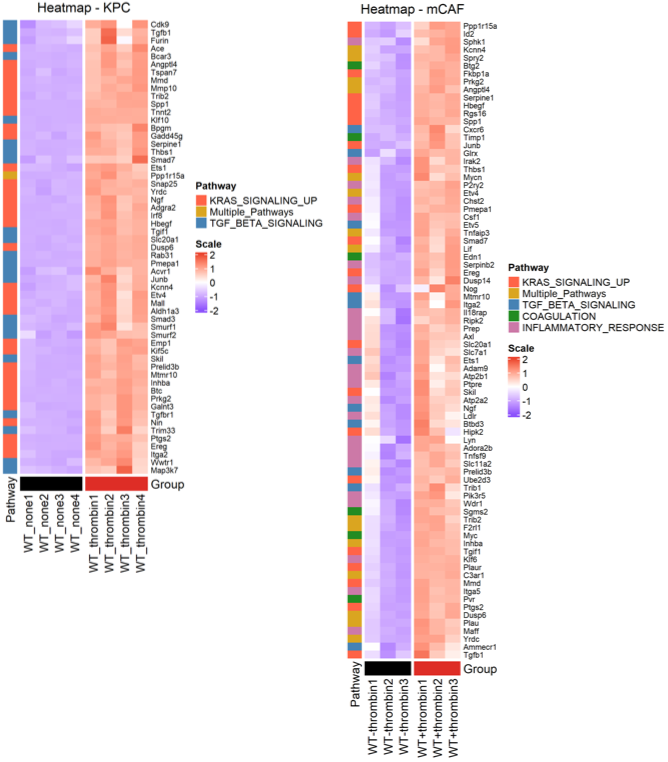

**Figure S3. Hallmark genes upregulated by thrombin-PAR1 signaling in KPC and mCAF cells.**

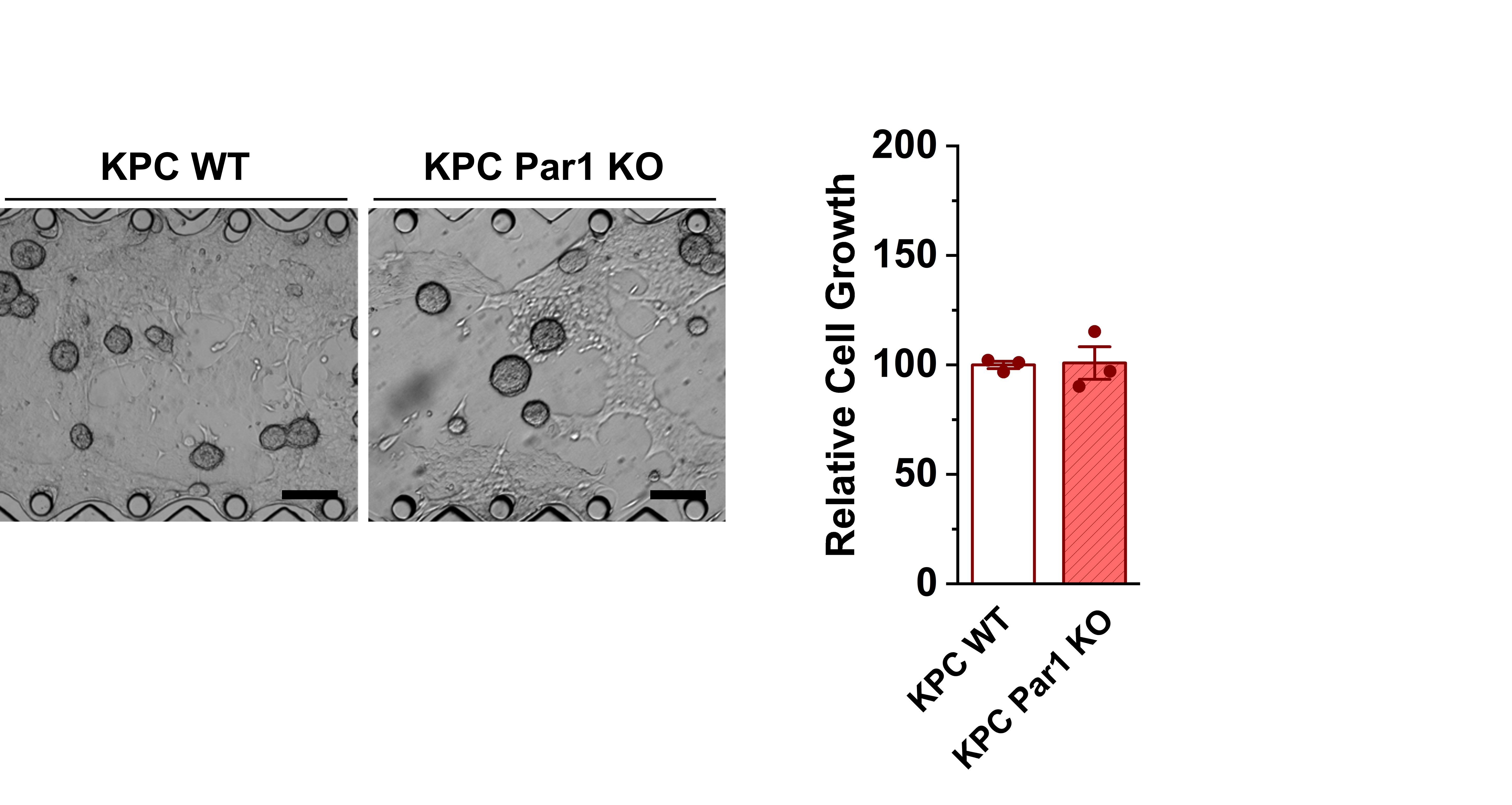

**Figure S4**. **KPC WT and KPC Par1 KO cell growth in monoculture**. Scale bar = 200 μm.

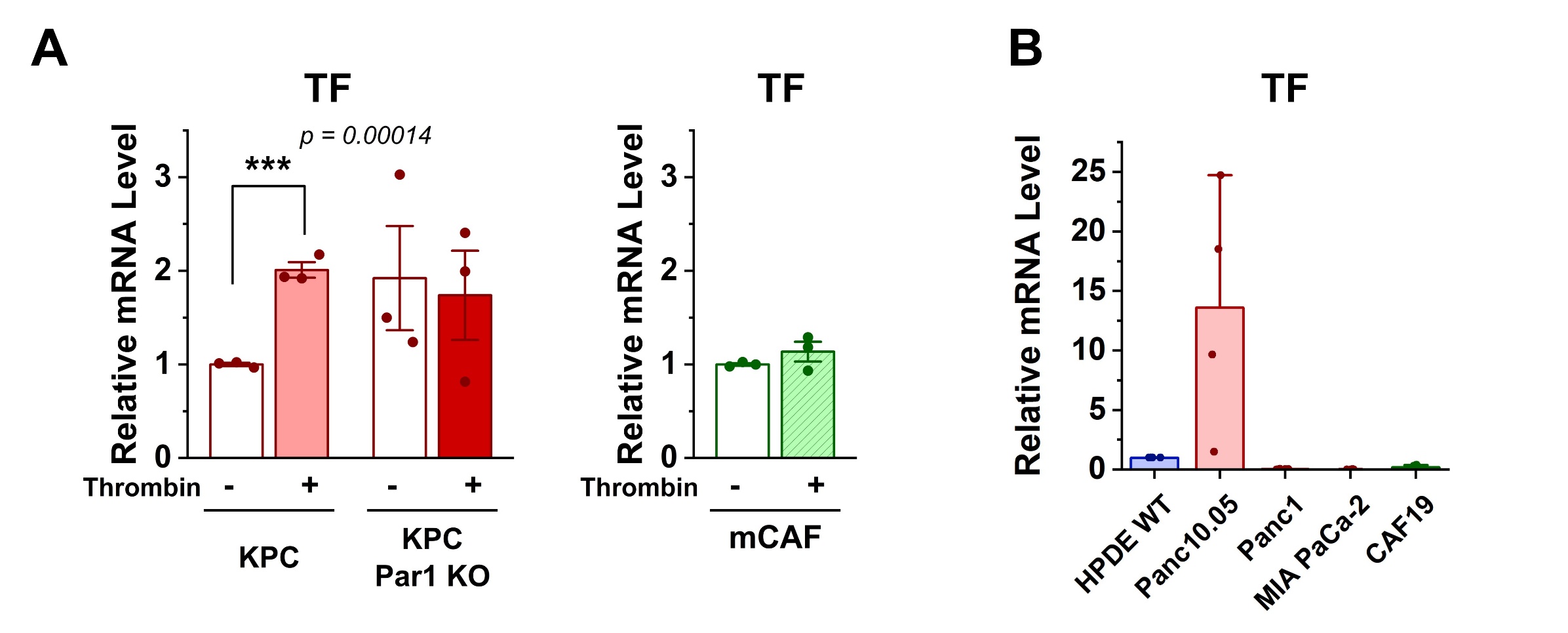

**Figure S5. Relative mRNA levels of tissue factor (TF) in human cell lines.**

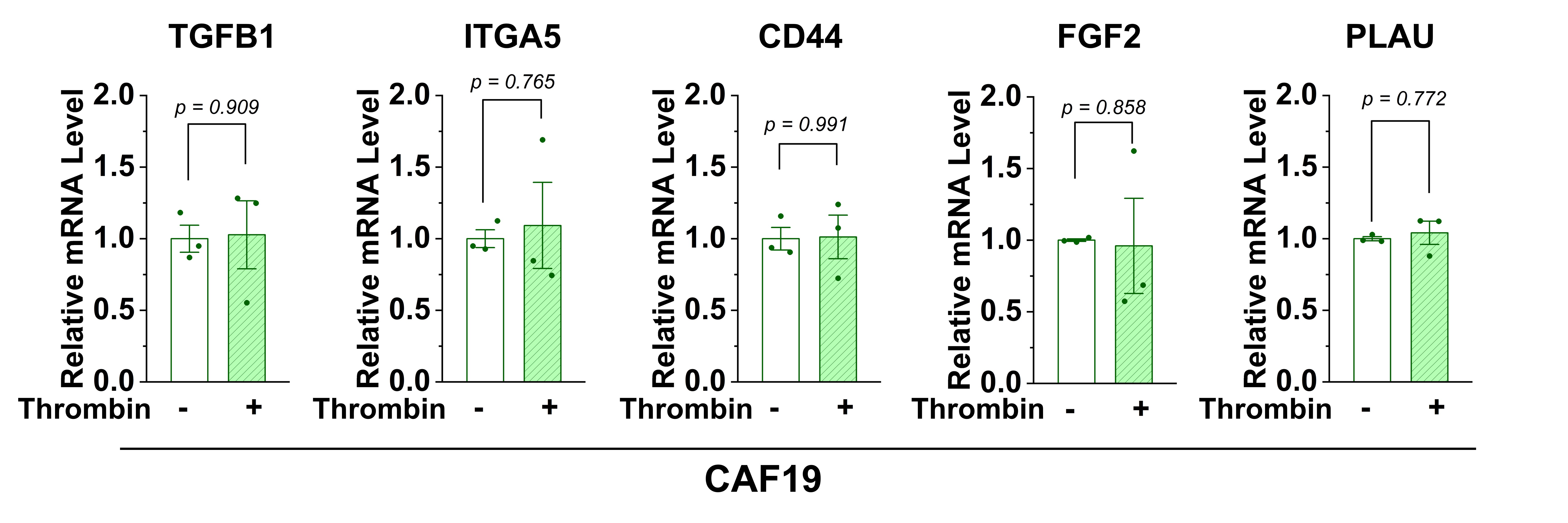

**Figure S6**. **Relative mRNA levels of fibrosis-related genes in CAF19 after thrombin stimulation.**
\

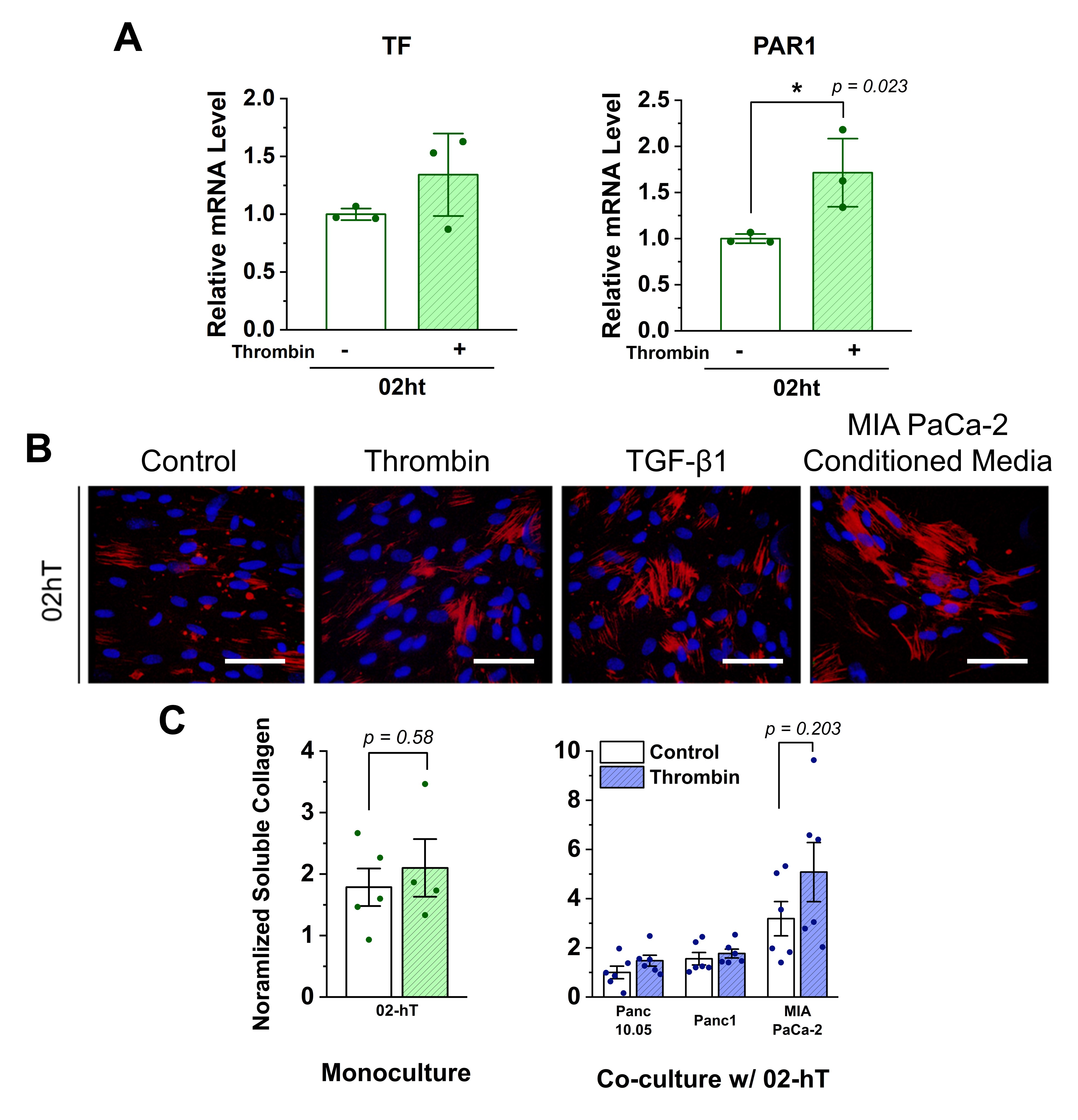

**Figure S7. Effect of thrombin treatment in a human CAF cell line 02hT.** (A) TF and PAR1 mRNA expressions. (B) αSMA expression measured by immunostaining. Scale bar = 100 µm; (C) Soluble collagen in monoculture and co-culture systems.

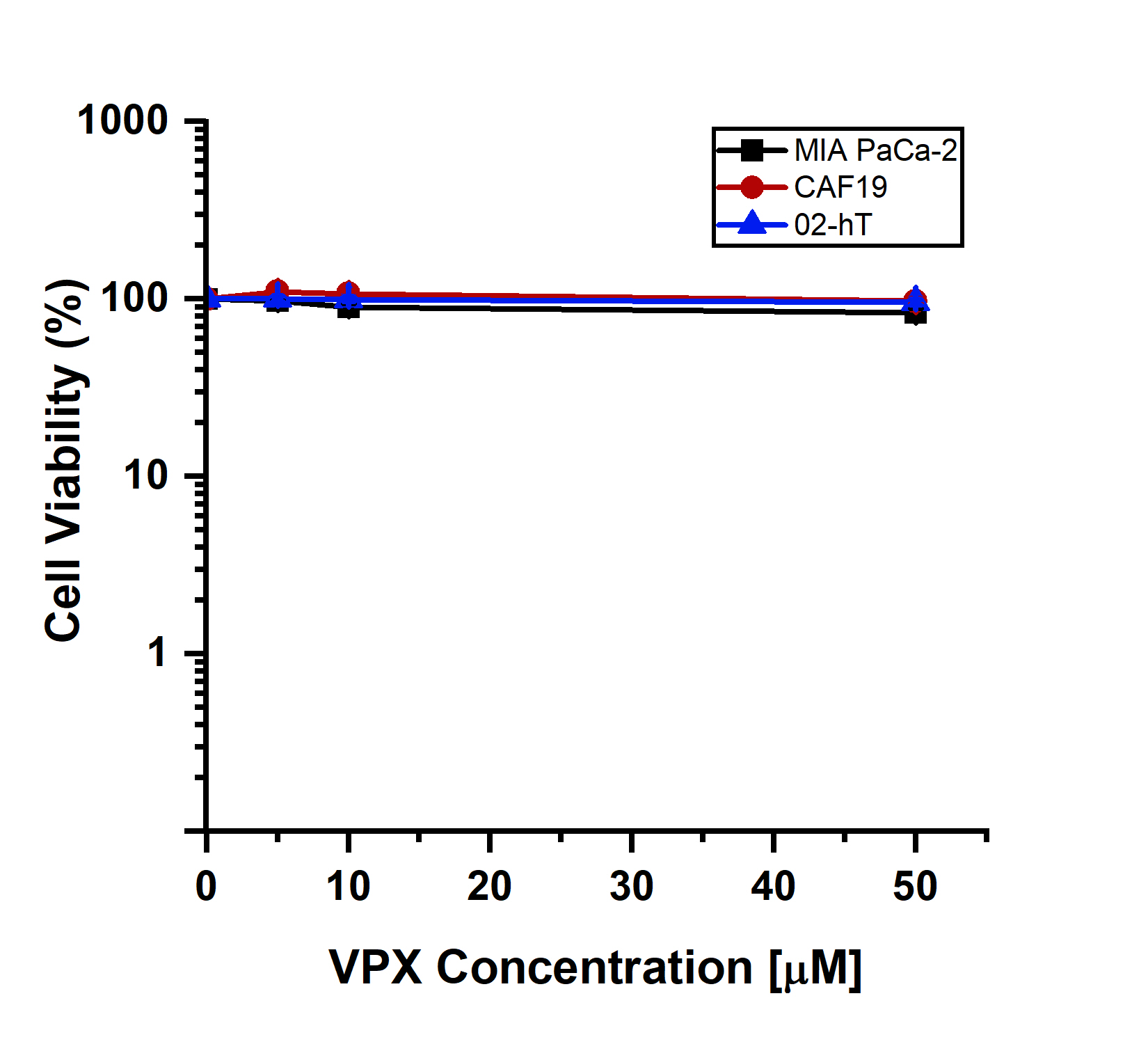

**Figure S8. Vorapaxar (VPX) cytotoxicity assay**. Cell viability was measured after 72-hour treatment with VPX, measured by alamarBlue.

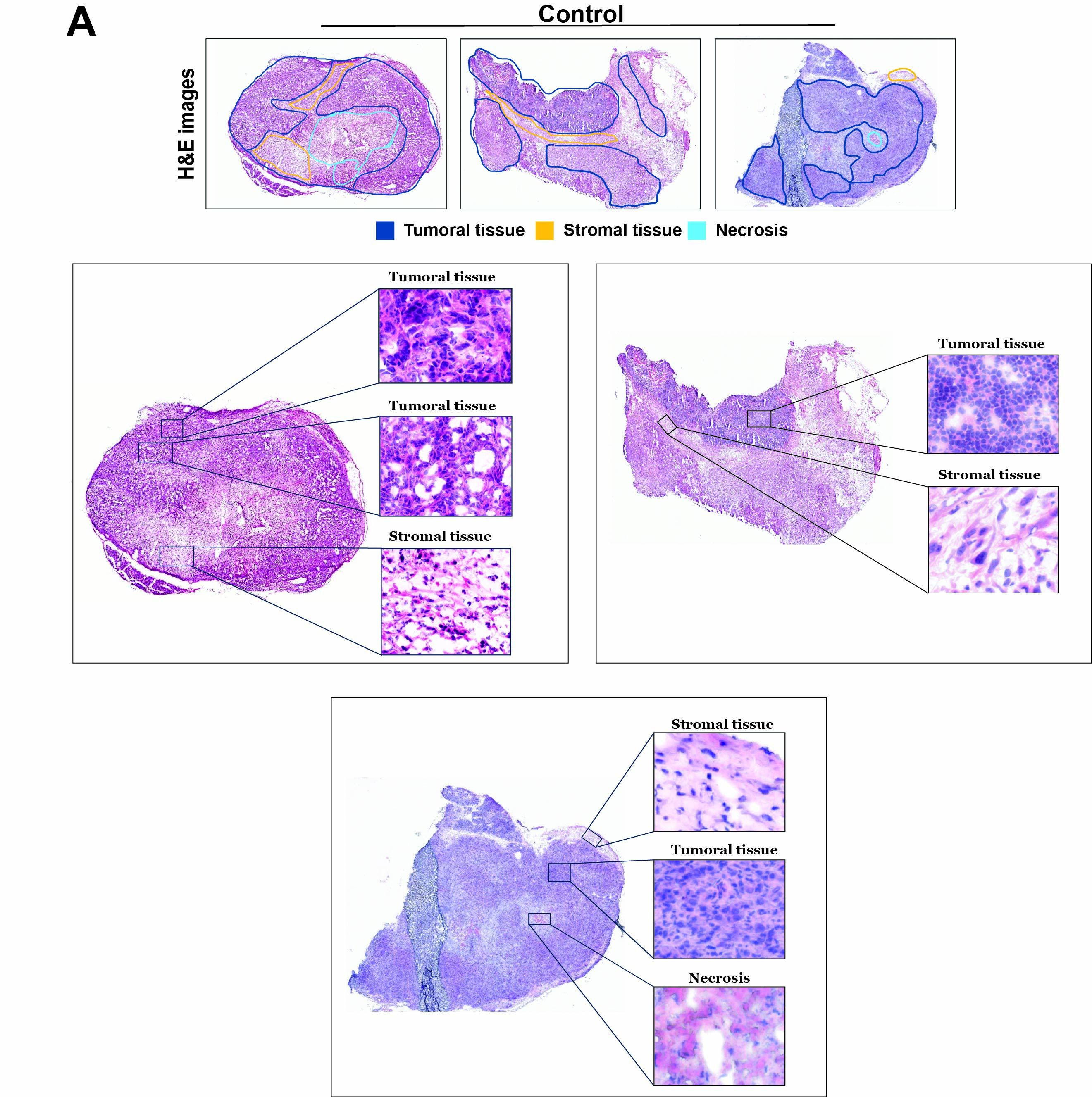

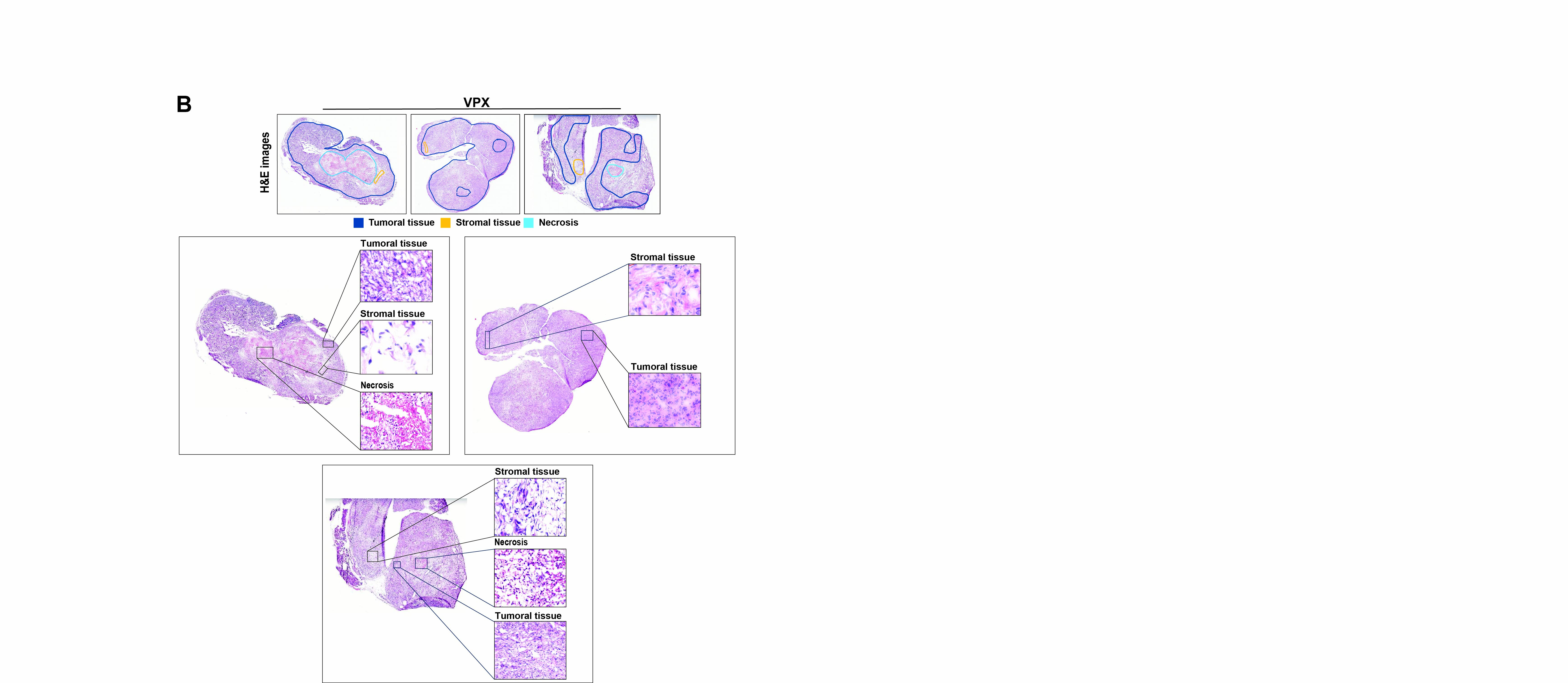

**Figure S9. Histology analysis of orthotopic tumors.** Tumor, stroma and necrotic tissue regions are annotated in control and VPX treatment groups. Because histopathologic assessment of cryosectioned tissues relies primarily on morphology and tissue context, it cannot provide fully definitive confirmation of cell identity, particularly for phenotypically overlapping or spatially heterogeneous cell populations.

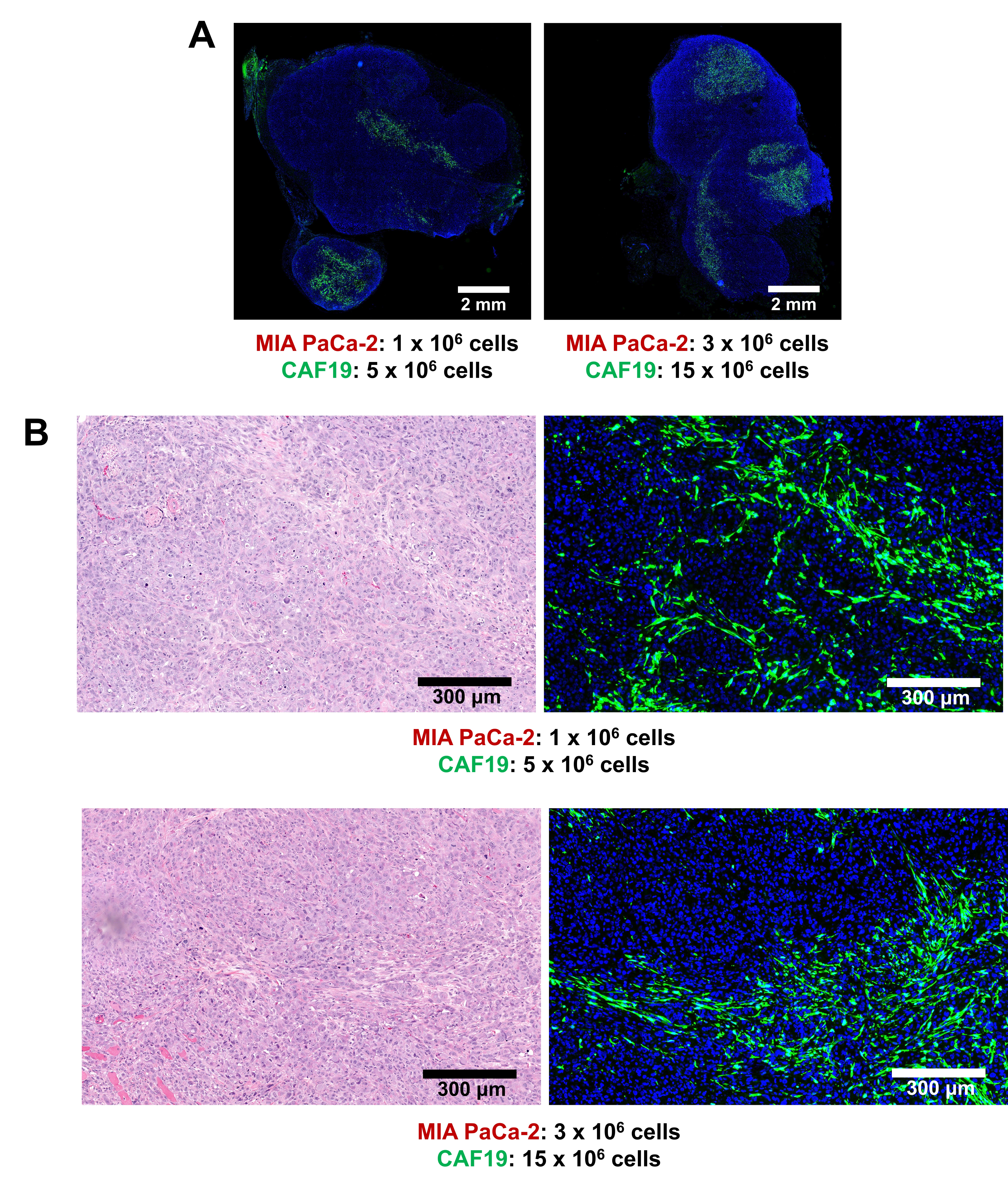

**Figure S10. Investigation of initial cell seeding concentration and final CAF population in xenograft model.** (A) Fluorescent images of whole tumor for two different cell numbers of PCc and CAF, while keeping the ratio consistent. (B) Magnified H&E and IF images of two cell numbers investigated. Significant FITC-CAF (anti-GFP) expression after 4 weeks of tumor growth validates 1:5 PCC:CAF ratio. Based on this results, 0.7 x 10^6^ : 3.5 x 10^6^ MIA PaCa-2: CAF19 cells per mice were inoculated, where the ratio is kept constant with reduced total cell numbers due to technical difficulties in providing large numbers of cells for the number of sample mice.
